# Glial involvement in mitochondrial metabolic alterations in Alzheimer’s disease

**DOI:** 10.64898/2026.11.20.768184

**Authors:** Kosei Hirata, Maiko Ono, Yuki Maeda, Yurika Abe, Rena Kono, Akiko Uematsu, Goh Kobayashi, Huseyin Erdogan, Kailash BP, Ken-ichi Dewa, Minami Motoyoshi, Jun Nagai, Hiroshi Shimizu, Akiyoshi Kakita, Towfique Raj, Yasuo Uchida, Yuhei Takado

**Author notes:** Corresponding author: Yuhei Takado. These authors contributed equally: Kosei Hirata and Maiko Ono.

## Abstract

Metabolic alterations in the posterior cingulate cortex (PCC) emerge early in Alzheimer’s disease (AD), but their molecular and cellular basis remains unclear. We hypothesized that impaired glial energy metabolism, particularly mitochondrial oxidative metabolism in astrocytes, underlies the metabolic vulnerability of the PCC in AD. To test this hypothesis, we integrated human PCC proteomics and transcriptomics with magnetic resonance spectroscopy in *App*^NL-G-F^ knock-in (APP-KI) mice. Human proteomic and transcriptomic analyses converged on reduced mitochondrial oxidative-metabolism signatures associated with AD pathology and cognition. APP-KI hippocampus showed proteomic and ^1^H-MRS alterations resembling those in the human AD PCC. Spatial transcriptomics identified widespread metabolic transcriptional changes and reduced mitochondrial pathway signatures in glia-enriched hippocampal clusters. Hyperpolarized [1-^13^C]pyruvate spectroscopy showed reduced bicarbonate labelling in APP-KI mice, consistent with impaired oxidative pyruvate metabolism. These findings reveal convergent mitochondrial alterations in human AD and APP-KI mice, implicating glial cells, particularly astrocytes, in the associated dysfunction.

---

Alzheimer’s disease (AD) is the leading cause of dementia worldwide and is characterized by progressive cognitive decline accompanied by the accumulation of amyloid-β plaques and tau neurofibrillary tangles^1^. Beyond these hallmark pathologies, converging evidence indicates that disturbances in brain energy metabolism represent a fundamental feature of AD^2^. Impaired cerebral glucose utilization is a well-established feature of AD, while alterations in glycolytic and mitochondrial metabolism have also been implicated in its pathophysiology^3–5^. These findings raise the possibility that impaired energy homeostasis contributes to neuronal dysfunction and cognitive decline.

The posterior cingulate cortex (PCC), a highly metabolically active hub of the default mode network, is particularly vulnerable to metabolic disturbances in AD^3,6^. The PCC is among the earliest regions to show metabolic abnormalities, with reduced fluorodeoxyglucose uptake on positron emission tomography (FDG-PET), consistent with impaired glucose utilization^7,8^. Longitudinal FDG-PET imaging in autosomal dominant AD indicates that reduced glucose utilization can precede regional atrophy and emerge years before estimated symptom onset^9^.

Neuron–astrocyte metabolic interactions play an important role in matching energy supply to neuronal activity^10^. Astrocytes contribute to brain energy homeostasis not only through glycolytic metabolism but also through mitochondrial oxidative metabolism, raising the possibility that astrocytic metabolic dysfunction contributes to the altered energy balance observed in AD. Beyond FDG-PET evidence of impaired glucose utilization, our previous proton magnetic resonance spectroscopy (^1^H-MRS) study demonstrated elevated lactate levels in the AD PCC that correlated with myo-inositol, an astrocyte-associated metabolite, suggesting a possible glial contribution to this metabolic phenotype^11^. These observations led us to hypothesize that impaired glial energy metabolism, particularly mitochondrial oxidative metabolism in astrocytes, contributes to metabolic dysfunction in AD. However, the molecular and cellular basis of these metabolic alterations remains poorly understood.

Addressing this question requires complementary approaches that capture different aspects of metabolic regulation. Recent transcriptomic studies, including single-cell and spatially resolved approaches, have revealed diverse cell-type-specific alterations in AD^12,13^, but transcript abundance alone cannot determine whether these changes translate into altered metabolic function. Proteomic profiling provides a complementary view by quantifying proteins involved in cellular metabolism, particularly because transcript and protein abundance do not always correspond in the AD brain^14^. However, protein abundance does not directly reflect enzymatic activity or metabolic flux. In vivo metabolic imaging provides complementary information by assessing metabolite levels and labelled-substrate conversion, offering insights into metabolic alterations that cannot be inferred from molecular profiles alone^15^. Together, these approaches offer a means to connect molecular and cellular alterations with in vivo metabolic dysfunction in AD.

Here, we integrated bulk proteomic and transcriptomic profiling of the human AD PCC with multimodal analyses of the hippocampus in *App*^NL-G-F^ knock-in (APP-KI) mice, including proteomics, spatial transcriptomics, ^1^H-MRS and hyperpolarized [1-^13^C]pyruvate MRS. Given the role of locus coeruleus (LC) noradrenergic signalling in regulating astrocytic function and metabolism^16–18^, we further examined associations between LC neuronal counts and PCC metabolic and glial protein abundance. Together, these approaches identified convergent alterations in mitochondrial energy metabolism across human AD and APP-KI mice, alongside in vivo evidence of reduced oxidative pyruvate metabolism and the involvement of glial cells, particularly astrocytes, in the metabolic phenotype.

## Results

### Mitochondrial energy-metabolism signatures are reduced in the AD PCC

We performed quantitative proteomic profiling of PCC samples from AD (*n* = 7) and control (*n* = 9) donors using mass spectrometry with sequential window acquisition of all theoretical mass spectra (SWATH-MS; Fig. 1a)^19^, quantifying 6,305 proteins, with gene set enrichment analysis (GSEA) as the primary analysis to assess coordinated pathway-level changes across the ranked proteome. Exploratory differential abundance analysis provided an overview of individual-protein changes (Fig. 1b). At nominal *P* < 0.05 and an absolute log_2_ fold change (|log_2_FC|) > 0.1, proteins with lower abundance in AD included complex I subunits (NDUFA6, NDUFS1), the complex III subunit UQCRC1, pyruvate dehydrogenase (PDH) complex components (PDHA1, PDHX) and the tricarboxylic acid (TCA) cycle enzyme SUCLA2, whereas those with higher abundance included glial and inflammatory markers, including GFAP, MAOB, GPNMB and HMOX1 (Supplementary Table 1).

**Fig. 1.**
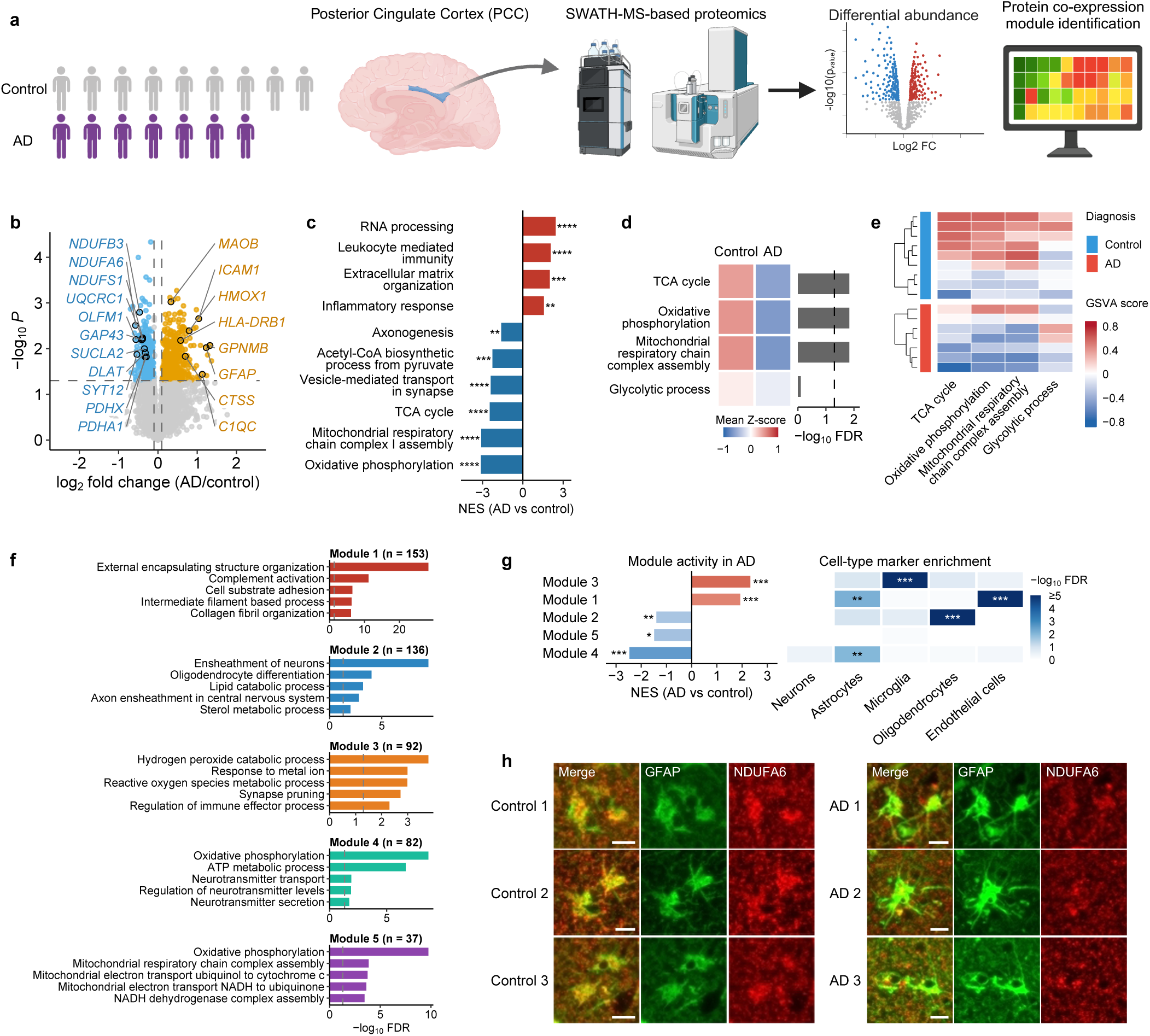
Proteomic profiling of the AD posterior cingulate cortex. (**a**) Study overview. Postmortem PCC tissue from AD (*n* = 7) and control (*n* = 9) cases was analysed by SWATH-MS-based quantitative proteomics, followed by differential abundance and protein co-expression analyses. (**b**) Volcano plot of differential protein abundance in AD versus control PCC. The y axis shows nominal −log_10_ P (two-sided Student’s *t*-test); dashed lines indicate *P* = 0.05 and |log_2_FC| = 0.1. Blue and orange points indicate proteins decreased and increased at nominal *P* < 0.05 and |log_2_FC| > 0.1, respectively. These results are exploratory; no individual protein reached proteome-wide FDR < 0.05. Labels indicate leading-edge proteins of the GSEA terms shown in c, together with GPNMB and MAOB; labelled points are outlined in black. Full results in Supplementary Table 1. (**c**) GSEA of GO BP terms across all quantified proteins ranked by the *t* statistic. Bars show normalized enrichment scores (NES; blue, lower in AD; red, higher in AD). Asterisks denote FDR: *FDR < 0.05, **FDR < 0.01, ***FDR < 0.001, ****FDR < 0.0001. Representative terms are shown; full results in Supplementary Table 2. (**d**) Group-mean GSVA scores for four energy-metabolism gene sets. Left, group means of GSVA scores z-scored across all 16 samples for each gene set (red, higher; blue, lower). Right, −log_10_ FDR from limma comparisons of AD and control samples; dashed line, FDR = 0.05. Full results in Supplementary Table 3. (**e**) Individual-sample GSVA heatmap for the same four gene sets. Rows represent samples and columns represent pathways; dendrograms indicate hierarchical clustering. The diagnosis annotation identifies control (blue) and AD (red) samples. Colours indicate GSVA enrichment scores, without z-scoring. (**f**) GO BP over-representation analysis of the five protein co-expression modules identified by CEMiTool. Bars show −log_10_ FDR and are coloured by module. Dashed lines indicate FDR = 0.05. Module sizes are shown in the titles. Representative terms are displayed; full results in Supplementary Table 4b. (**g**) Module enrichment in AD and cell-type marker enrichment. Modules are ordered by NES. Left, NES from CEMiTool gene set enrichment analysis (red, higher in AD; blue, lower in AD). Right, enrichment for curated brain cell-type marker sets^4^, assessed by one-sided Fisher’s exact test with BH correction within each cell type. Colours indicate −log_10_ FDR, with values ≥5 shown at the top of the scale. Asterisks denote FDR: *FDR < 0.05, **FDR < 0.01, ***FDR < 0.001. Full results in Supplementary Table 4c,d. (**h**) Double immunofluorescence images of GFAP (green) and NDUFA6 (red) in human PCC from three control (left) and three AD cases (right) included in the proteomic cohort. Each row shows one case, with merged, GFAP and NDUFA6 channels from left to right within each group. Row numbers do not indicate matched case pairs. Sections were obtained from the hemisphere contralateral to the tissue used for proteomics. NDUFA6 immunoreactivity overlaps with GFAP-positive profiles. Images are qualitative and were not used to quantify diagnostic-group differences. Scale bars, 10 μm. Lower-magnification images are shown in Supplementary Fig. 1. Abbreviations: AD, Alzheimer’s disease; PCC, posterior cingulate cortex; SWATH-MS, mass spectrometry with sequential window acquisition of all theoretical mass spectra; GSEA, gene set enrichment analysis; GO BP, Gene Ontology Biological Process; FDR, false discovery rate; GSVA, gene set variation analysis; BH, Benjamini–Hochberg; GFAP, glial fibrillary acidic protein; log_2_FC, log_2_ fold change.

GSEA across all quantified proteins identified 545 Gene Ontology Biological Process (GO BP) terms at a false discovery rate (FDR) < 0.05 (Supplementary Table 2). Negatively enriched terms were dominated by mitochondrial energy metabolism, including oxidative phosphorylation (normalized enrichment score, NES = −3.12, FDR = 4.8 × 10^−17^), mitochondrial respiratory chain complex I assembly (NES = −3.09, FDR = 2.6 × 10^−13^), the TCA cycle (NES = −2.46, FDR = 2.3 × 10^−5^) and acetyl-CoA biosynthesis from pyruvate (NES = −2.26, FDR = 3.5 × 10^−4^). Synaptic vesicle transport was also reduced, whereas RNA processing, immune and inflammatory processes, and extracellular matrix organization were positively enriched (Fig. 1c).

Gene set variation analysis (GSVA) using four selected energy-metabolism gene sets showed lower scores for mitochondrial respiratory chain complex assembly, oxidative phosphorylation and the TCA cycle (all FDR = 0.015), with no significant difference in glycolysis (FDR = 0.79; Fig. 1d,e; Supplementary Table 3).

To identify networks of proteins with correlated abundance across donors, we used CEMiTool, which identified five co-expression modules (M1–M5; 37–153 proteins). Among these, M4 and M5 were enriched for energy-metabolism-related processes and were both suppressed in AD (Fig. 1f,g; Supplementary Table 4a–e). M4 (82 proteins) showed the strongest suppression (NES = −2.47, FDR = 5.1 × 10^−8^) and was enriched for astrocyte markers (FDR = 9.8 × 10^−3^). It contained respiratory-chain subunits (NDUFA5, NDUFA6, COX7A1, SDHC), glutamine synthetase (GLUL) and synaptic proteins (SYP, RPH3A, SNCA), linking mitochondrial and neurotransmitter-related proteins within the same module. This organization suggests coordinated variation in energy metabolism and processes supporting astrocyte–neuron interactions. M5 (37 proteins) was also suppressed (NES = −1.49, FDR = 0.035) and was enriched for respiratory-chain components, without significant enrichment for the tested cell-type marker sets. The remaining modules captured myelin-associated processes (M2), immune and microglial processes (M3), and extracellular matrix and complement processes with endothelial and astrocyte marker enrichment (M1). M2 was suppressed in AD, whereas M1 and M3 were increased (Fig. 1g; Supplementary Table 4c,d).

To examine the cellular localization of a mitochondrial protein implicated by both differential abundance and network analyses, we selected NDUFA6, a respiratory-chain complex I accessory subunit that showed reduced abundance in the exploratory proteomic analysis and belonged to the downregulated, astrocyte-marker-enriched M4 module. Double immunofluorescence staining for NDUFA6 and GFAP was performed on PCC sections from three control and three AD donors in the proteomic cohort, using tissue from the hemisphere contralateral to that sampled for proteomics. NDUFA6 immunoreactivity overlapped with GFAP-positive astrocytic profiles in both groups, supporting the presence of NDUFA6 in astrocytes (Fig. 1h; Supplementary Fig. 1).

### Independent PCC transcriptomics corroborates reduced mitochondrial pathway signatures

We analysed PCC bulk RNA sequencing (RNA-seq) from the Religious Orders Study and Rush Memory and Aging Project (ROSMAP)^20^, comprising 117 pathologically defined AD and 69 cases with no cognitive impairment (NCI) (Table 1), identifying 991 decreased and 1,110 increased genes at FDR < 0.05 and |log_2_FC| > 0.1 (Fig. 2a; Supplementary Table 5). Decreased transcripts included the complex I subunits *NDUFA6* and *NDUFS1*, the PDH-complex component *PDHA1* and the TCA-cycle enzyme *SUCLA2*, consistent with the proteomic findings. GSEA identified 323 significant GO BP terms (Supplementary Table 6). Mitochondrial pathways were concordantly decreased across the proteomic and transcriptomic datasets, including mitochondrial translation (NES = −2.64, FDR = 1.8 × 10^−11^), the TCA cycle (NES = −2.54, FDR = 1.3 × 10^−6^), oxidative phosphorylation (NES = −2.43, FDR = 2.5 × 10^−9^), respiratory chain complex assembly and acetyl-CoA biosynthesis from pyruvate (Fig. 2b). Ribosome biogenesis and rRNA processing were also reduced. Increased pathways involved developmental and cell-fate programmes, glial differentiation and insulin receptor signalling, with leading-edge genes including *WNT6*, *SFRP2* and *NOG* (canonical Wnt signalling), *CXCR4* (glial cell differentiation) and *FOXO4* (insulin receptor signalling).

**Fig. 2.**
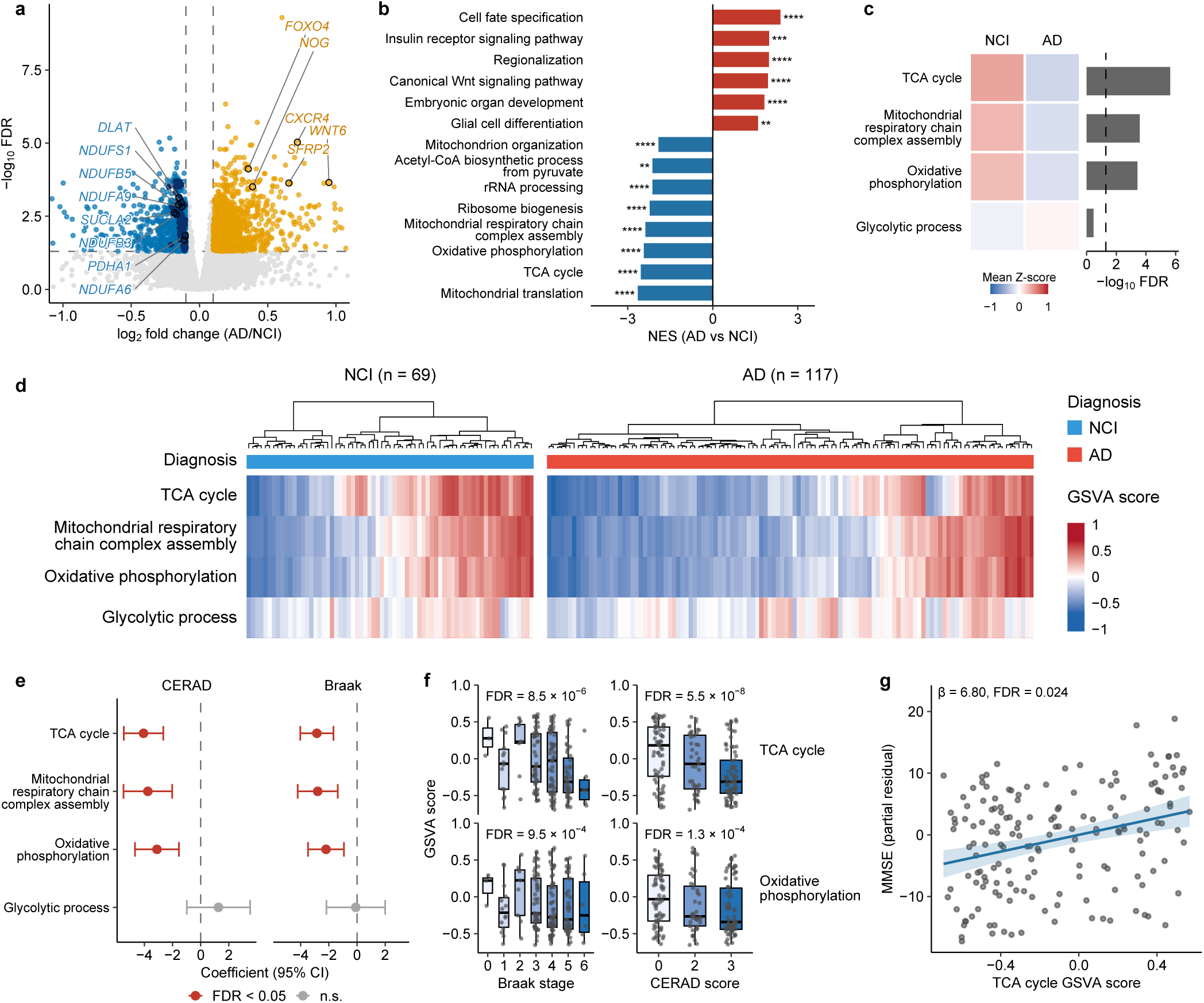
Transcriptomic profiling of the human AD PCC reveals mitochondrial metabolic alterations associated with neuropathology and cognition. **(a)** Differential gene expression in PCC samples from pathologically defined AD cases (*n* = 117) and controls with no cognitive impairment (NCI; *n* = 69) in ROSMAP. Each point represents a gene. Orange and blue indicate increased and decreased expression in AD, respectively, at FDR < 0.05 and |log_2_ fold change| > 0.1; grey indicates genes not meeting these criteria. Dark blue highlights decreased genes belonging to selected energy-metabolism pathways. Labels indicate leading-edge genes of the GSEA terms shown in b. Dashed lines indicate the significance and fold-change thresholds. Differential expression was assessed using DESeq2 with Benjamini–Hochberg correction. **(b)** Gene set enrichment analysis (GSEA) of Gene Ontology Biological Process (GO BP) terms, with genes ranked by the DESeq2 Wald statistic. Bars show normalized enrichment scores (NES; red, enriched among increased genes; blue, enriched among decreased genes). Representative terms are displayed. Asterisks indicate FDR: *FDR < 0.05, **FDR < 0.01, ***FDR < 0.001 and ****FDR < 0.0001. **(c)** Group comparison of gene set variation analysis (GSVA) scores for four energy-metabolism pathways. Left, group means of pathway-wise z-scored GSVA scores after covariate adjustment for visualization (red, higher; blue, lower). Right, −log_10_ false discovery rate (FDR) from covariate-adjusted limma models; the dashed line indicates FDR = 0.05. Models included age at death, postmortem interval (PMI), RNA integrity number (RIN), sequencing batch and sex. **(d)** Individual-sample GSVA heatmap for the same four pathways. Columns represent donors and rows represent pathways. Samples were hierarchically clustered within each diagnostic group. Colours indicate GSVA scores without z-scoring; the annotation bar indicates diagnosis. **(e)** Associations between GSVA pathway scores and CERAD neuritic plaque score or Braak neurofibrillary tangle stage across the combined AD and NCI groups (*n* = 186). Points and horizontal bars show regression coefficients and 95% confidence intervals from proportional-odds ordinal logistic regression, with neuropathological score as the outcome and pathway score as the predictor, adjusted for the covariates listed in c. Red indicates FDR < 0.05; grey indicates non-significant associations. BH correction was applied separately within each neuropathological trait. **(f)** TCA-cycle and oxidative phosphorylation GSVA scores stratified by Braak stage or CERAD score in the same cohort as e. Points represent individual donors. Boxes show the median and interquartile range; whiskers extend to the most extreme values within 1.5 times the interquartile range. Displayed FDR values derive from the covariate-adjusted ordinal regression models in e, rather than pairwise comparisons between categories. **(g)** Association between TCA-cycle GSVA score and antemortem Mini-Mental State Examination (MMSE) score within clinically diagnosed AD cases (*n* = 165). Points represent individual donors; the vertical axis shows MMSE partial residuals from a linear model adjusted for the covariates listed in c. The line and shaded band indicate the fitted relationship and its 95% confidence interval. The regression coefficient and FDR are shown; BH correction was applied across the two tested pathways, TCA cycle and oxidative phosphorylation. Abbreviations: AD, Alzheimer’s disease; PCC, posterior cingulate cortex; ROSMAP, Religious Orders Study and Rush Memory and Aging Project; NCI, no cognitive impairment; CERAD, Consortium to Establish a Registry for Alzheimer’s Disease; TCA, tricarboxylic acid; CI, confidence interval; n.s., not significant.

GSVA using the same four energy-metabolism gene sets as in the proteomic analysis showed lower scores for all three mitochondrial pathways in AD, with the strongest evidence for the TCA cycle (FDR = 2.3 × 10^−6^), and no significant difference in glycolysis (FDR = 0.34; Fig. 2c; Supplementary Table 7a). Individual scores varied within both groups (Fig. 2d). Across the combined AD and NCI groups, mitochondrial scores were negatively associated with the Consortium to Establish a Registry for Alzheimer’s Disease (CERAD) score and Braak stage, most strongly for the TCA cycle (CERAD FDR = 5.5 × 10^−8^; Braak FDR = 8.5 × 10^−6^; Fig. 2e,f; Supplementary Table 7b). TCA-cycle scores were positively associated with antemortem Mini-Mental State Examination (MMSE) scores (β = 6.80, FDR = 0.024), whereas oxidative phosphorylation scores were not (FDR = 0.36; Fig. 2g; Supplementary Table 7c). Thus, the mitochondrial transcriptomic signature was associated with AD neuropathology, and the TCA-cycle score additionally tracked cognitive performance within clinical AD.

CEMiTool identified 11 RNA co-expression modules spanning synaptic and neuronal, immune, vascular and extracellular-matrix, astrocytic, myelination, proteostatic and interferon-response functions, but no energy-metabolism module (Supplementary Fig. 2; Supplementary Table 8a– e); only 2 of the 149 genes annotated to oxidative phosphorylation (GO:0006119) passed CEMiTool’s variance-based filter. The neuronal marker-enriched modules M1 and M2 and module M8 were positively associated with MMSE (adjusted β = 0.24–0.27, FDR ≤ 6.2 × 10^−3^), and M2 and M8 were decreased in AD (NES = −1.72 and −2.62). M8 showed no significant pathway enrichment but included parvalbumin-interneuron genes (PVALB, KCNC1 and SCN1B). Conversely, the myelination module M6, the protein-folding and heat-shock module M9 and the astrocyte marker-enriched module M5 were increased in AD (NES = 2.76, 2.05 and 1.51; Supplementary Table 8c).

### The APP-KI hippocampal proteome shares the human mitochondrial signature

Quantitative proteomics of 18-month-old APP-KI and wild-type (WT) hippocampus (*n* = 6 per group; 5,700 proteins) identified 249 increased and 99 decreased proteins at FDR < 0.05 (Fig. 3a; Supplementary Table 9a). GSEA revealed reduced enrichment of oxidative phosphorylation (NES = −1.98, FDR = 0.0013) and the TCA cycle (NES = −1.94, FDR = 0.016; Fig. 3b; Supplementary Table 9b).

**Fig. 3.**
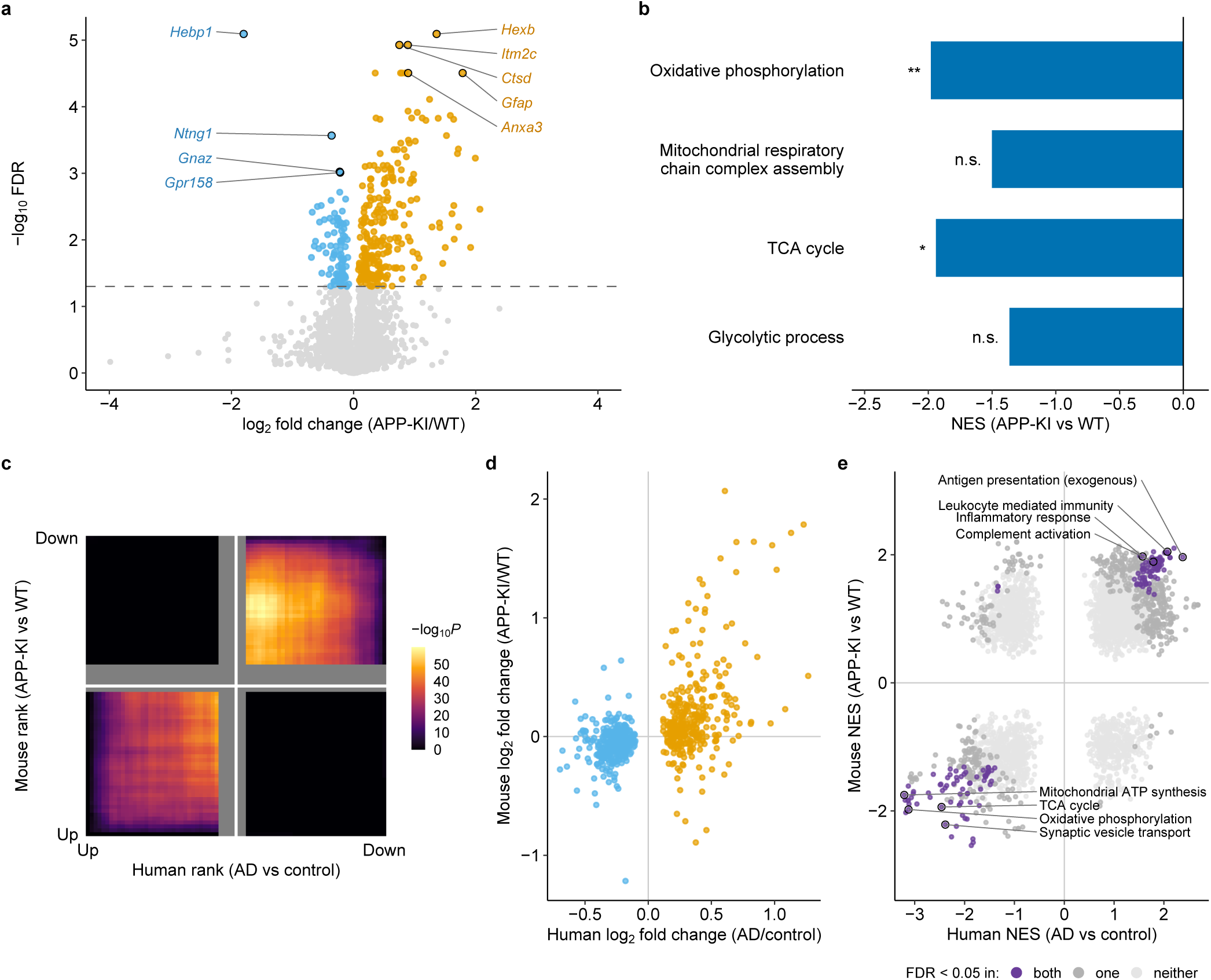
The APP-KI hippocampal proteome shares mitochondrial metabolic alterations with the human AD PCC. **(a)** Differential protein abundance in hippocampal samples from 18-month-old APP-KI and wild-type (WT) mice (*n* = 6 per group). Each point represents a protein. Orange and blue indicate higher and lower abundance in APP-KI mice, respectively, at FDR < 0.05; grey indicates non-significant differences. The dashed line indicates FDR = 0.05. Selected proteins are labelled by their corresponding gene symbols. Differential abundance was assessed using two-sided Student’s *t*-tests with BH correction across all quantified proteins. **(b)** GSEA of four energy-metabolism GO BP gene sets, with all quantified proteins ranked by the *t* statistic. Bars show NES; negative values indicate enrichment among proteins with lower abundance in APP-KI mice. Asterisks indicate FDR: *FDR < 0.05 and **FDR < 0.01; n.s., not significant. **(c)** Rank–rank hypergeometric overlap analysis (RRHO2) across 4,881 one-to-one human– mouse ortholog pairs quantified in both proteomic datasets. Proteins were ranked independently in each species by signed −log_10_ *P*. Colours indicate −log_10_ *P* from hypergeometric overlap tests. The lower-left and upper-right quadrants represent concordantly increased and decreased proteins, respectively; the remaining quadrants represent changes in opposite directions. **(d)** Comparison of human and mouse log_2_ fold changes for 700 ortholog pairs meeting the exploratory human differential-abundance threshold of nominal *P* < 0.05 and |log_2_ fold change| > 0.1. Each point represents an ortholog pair; orange and blue indicate higher and lower abundance in human AD, respectively. Changes were concordant in direction for 510 of 700 pairs (72.9%; two-sided binomial *P* = 1.1 × 10^−34^ against a null proportion of 0.5; Spearman ρ = 0.52). **(e)** Comparison of GSEA NES across 3,717 GO BP terms tested in both species. Each point represents a term. Purple indicates FDR < 0.05 in both species, dark grey in one species and light grey in neither species. Selected terms are labelled using abbreviated names. NES were positively correlated across species (Spearman ρ = 0.46), and 152 of the 155 terms significant in both species changed in the same direction. GO BP terms share genes and are not statistically independent. Abbreviations: APP-KI, *App*^NL-G-F^ knock-in; AD, Alzheimer’s disease; PCC, posterior cingulate cortex; WT, wild type; BH, Benjamini–Hochberg; FDR, false discovery rate; GSEA, gene set enrichment analysis; GO BP, Gene Ontology Biological Process; NES, normalized enrichment score; TCA, tricarboxylic acid; n.s., not significant.

Comparison of protein abundance changes across 4,881 one-to-one human–mouse ortholog pairs revealed concordance between human AD PCC and APP-KI hippocampus. Rank–rank hypergeometric overlap analysis (RRHO2) detected overlap in both concordant quadrants, with little overlap in discordant quadrants (Fig. 3c). Of 700 orthologs meeting the exploratory human threshold, 510 (72.9%) changed in the same direction in mice (binomial *P* = 1.1 × 10^−34^; Spearman ρ = 0.52; Fig. 3d; Supplementary Table 10a). NES from GSEA performed separately in each species were positively correlated across the 3,717 GO BP terms tested in both (Spearman ρ = 0.46). Of the 155 terms reaching FDR < 0.05 in both species, 152 changed concordantly, including mitochondrial and respiratory-chain terms and synaptic terms decreased in both species and immune terms increased in both (Fig. 3e; Supplementary Table 10b).

### APP-KI mice show lactate and *myo*-inositol changes resembling those in human AD PCC

To assess the in vivo metabolic phenotype, we performed ^1^H-MRS in APP-KI hippocampus and cortex (Fig. 4a,b), comparing the findings with our human AD PCC study^11^. APP-KI mice were examined at 3, 6, 8, 10 and 13 months; WT comparators were studied at 6 and 12 months. Hippocampal lactate was elevated at 3 months (*P* = 0.0004), 10 months (*P* = 0.0248) and 13 months (*P* = 0.0052), while *myo*-inositol was increased at 8, 10 and 13 months (*P* = 0.0398, 0.0117 and < 0.0001, respectively). Total N-acetylaspartate (tNAA) and total choline (tCho) did not differ significantly (Fig. 4c). Hippocampal glutamine was elevated at 8, 10 and 13 months (*P* = 0.0007, 0.0053 and 0.0207, respectively), over the same age range as the myo-inositol increase; other metabolites are shown in Supplementary Fig. 3. The absence of same-age WT controls at 3 months limits inference about the onset of these changes.

**Fig. 4.**
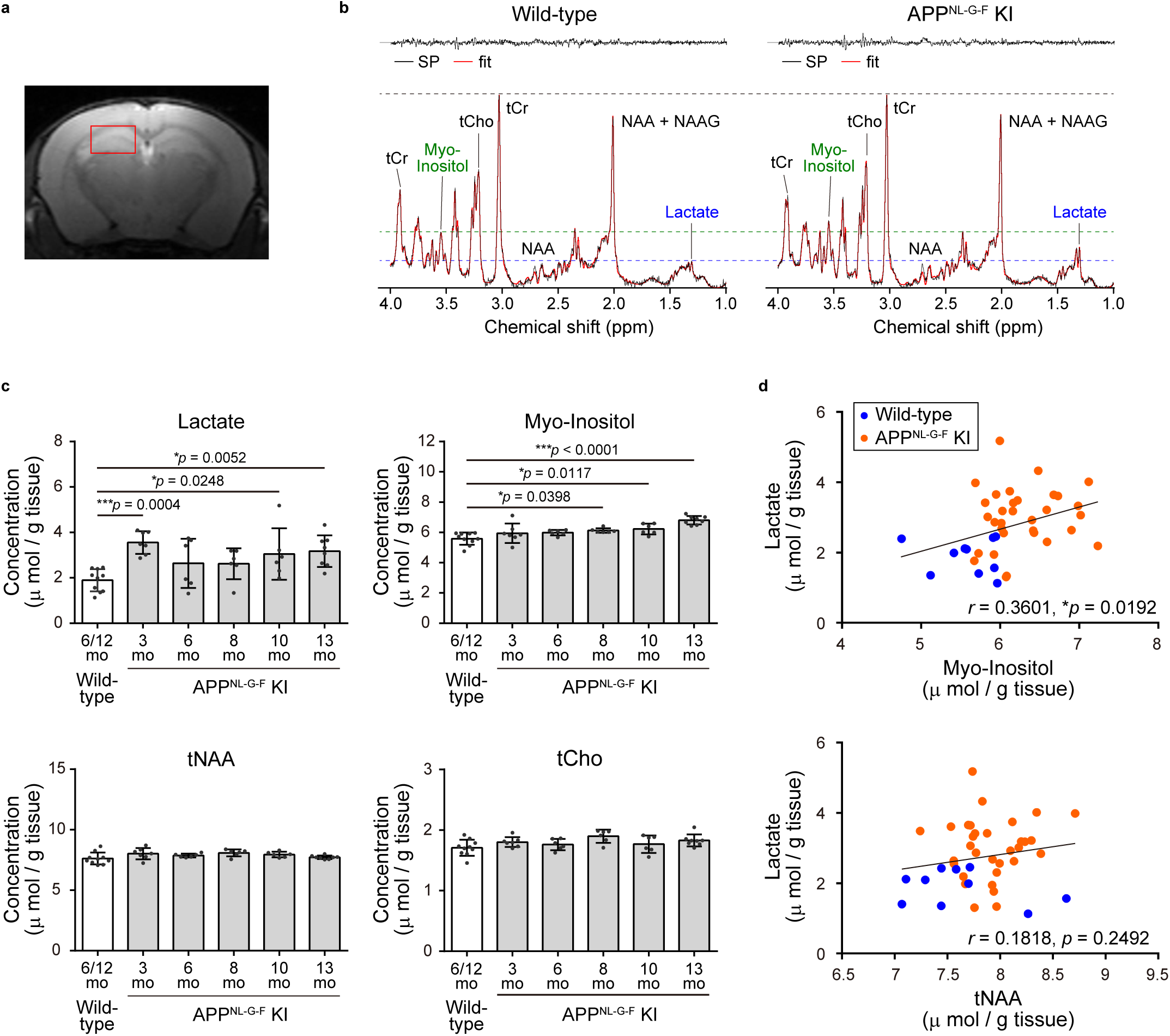
Magnetic resonance spectroscopy in APP-KI mice. Magnetic resonance spectroscopy (MRS)-based metabolite assessment in the hippocampus of APP-KI and wild-type mice. (**a**) Representative single-voxel placement (red squares) in the hippocampus (1.2 × 2.0 × 2.2 mm^3^) overlaid on a coronal T_2_-weighted MR image. (**b**) Representative single-voxel ^1^H-MRS (PRESS) spectra and corresponding fits from 13-month-old wild-type and APP-KI mice. tCr = creatine + phosphocreatine, tCho = glycerophosphocholine + phosphocholine, NAA = N-acetylaspartate, NAAG = N-acetylaspartylglutamate. (**c**) Hippocampal concentrations of lactate, *myo*-inositol, tNAA, and tCho in wild-type mice (6 and 12 months) and APP-KI mice (3, 6, 8, 10, and 13 months). Data are shown as individual values with mean ± standard deviation. \**P* < 0.05, \*\*\**P* < 0.0005 by one-way ANOVA followed by Dunnett’s multiple comparisons test. (**d**) Correlations between hippocampal metabolite concentrations, specifically lactate and *myo*-inositol (upper), and lactate and tNAA (lower), across all animals. \**P* < 0.05 using Pearson’s correlation analysis. Abbreviations: APP-KI, *App*^NL-G-F^ knock-in; MRS, magnetic resonance spectroscopy; ^1^H-MRS, proton magnetic resonance spectroscopy; MR, magnetic resonance; PRESS, point-resolved spectroscopy; tNAA, total N-acetylaspartate (NAA + NAAG); tCho, total choline; ANOVA, analysis of variance.

Across the pooled animals, hippocampal lactate correlated with *myo*-inositol (*r* = 0.36, *P* = 0.0192) but not tNAA (*r* = 0.18, *P* = 0.2492; Fig. 4d). The cortex also showed elevated lactate and *myo*-inositol and a positive correlation between them (*r* = 0.44, *P* = 0.0033; Supplementary Fig. 4). These observations resembled the human PCC findings, although the pooled correlations do not separate age and genotype effects.

In rTg4510 mice, lactate showed no significant change across the examined age groups, while *myo*-inositol was transiently reduced at 3–4 months (*P* = 0.0482). tNAA was reduced at 5–6 and 7–8 months (both *P* < 0.0001), and lactate did not correlate with *myo*-inositol (*r* = 0.013, *P* = 0.866; Supplementary Fig. 5). The APP-KI profile therefore more closely resembled the human AD PCC findings. Together, these proteomic and ^1^H-MRS similarities support further investigation of the cellular and metabolic alterations in the APP-KI hippocampus.

### Glia-enriched spatial clusters show reduced mitochondrial pathway signatures in the APP-KI hippocampus

To investigate the regional and cellular context of the MRS phenotype, we performed spatial transcriptomic profiling on brain tissue collected from the same APP-KI and WT mice after in vivo MRS at 6 months of age. Visium v2 profiling of whole-hemisphere sections delineated 12 anatomical regions (*n* = 4 per group; Fig. 5a). Disease-associated astrocyte (DAA) and disease-associated microglia (DAM) signature scores were each increased in 10 of the 12 regions, with the largest effects in cortical regions (Fig. 5b,c; Supplementary Fig. 6; Supplementary Table 11c). Energy-metabolism scores revealed widespread increases in glycolysis-related transcriptional signatures and decreases in mitochondrial pathway signatures across multiple brain regions: glycolysis scores increased in 9 regions, whereas oxidative phosphorylation and TCA-cycle scores decreased in 7 and 5 regions, respectively (FDR < 0.05; Fig. 5b,c; Supplementary Table 11c). In the hippocampus, glycolysis scores were increased (FDR = 0.0011) and oxidative phosphorylation scores were decreased (FDR = 0.036), whereas the decrease in TCA-cycle scores did not reach significance (FDR = 0.084).

**Fig. 5.**
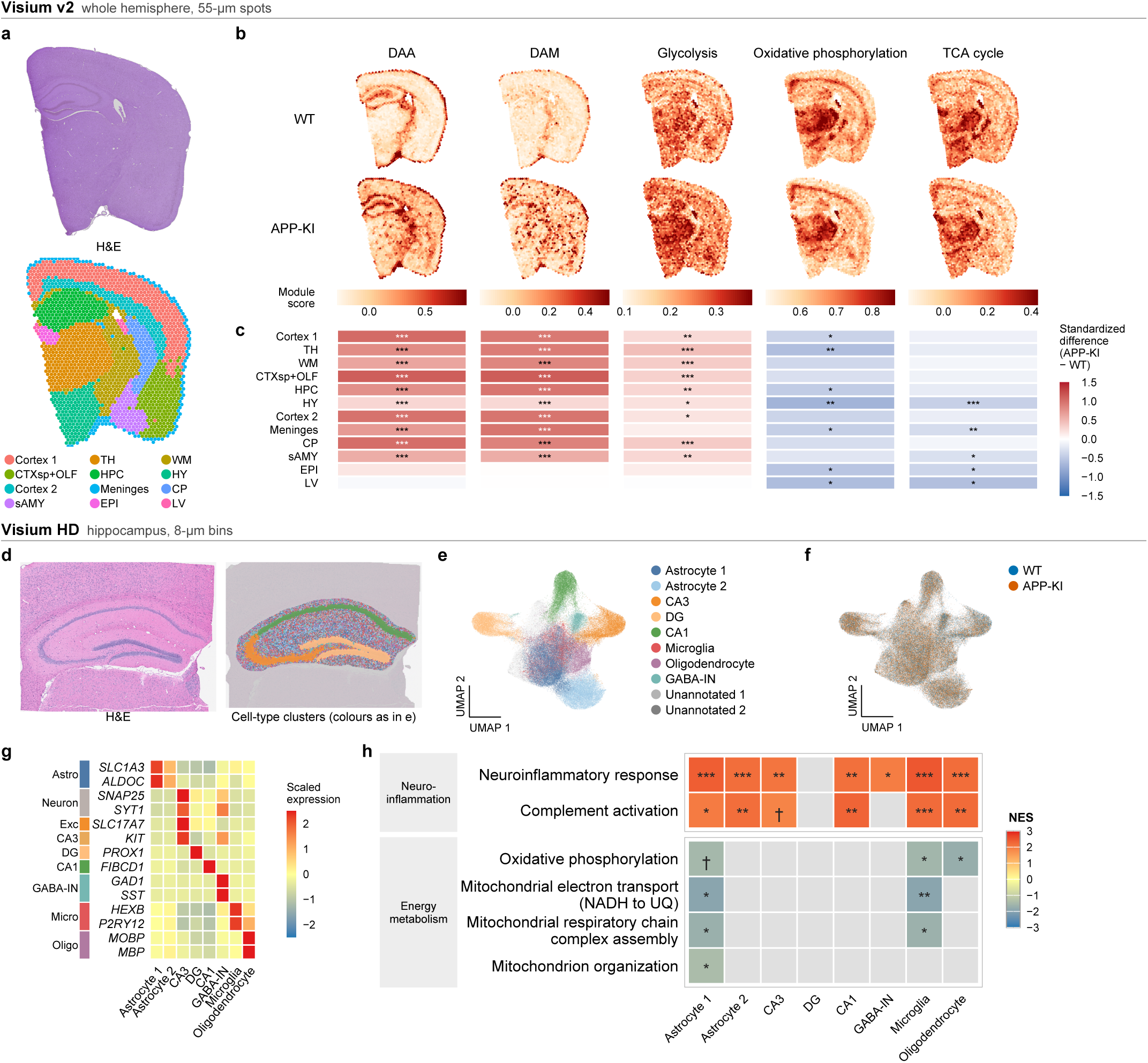
Spatial transcriptomic profiling reveals glial activation and altered energy-metabolism signatures in APP-KI mice. (a–c) Visium v2 profiling of whole hemispheres (55-µm spots). (d–h) Visium HD profiling of the hippocampus (8-µm bins). **(a)** Representative haematoxylin and eosin (H&E)-stained section (top) and spatial clustering map (bottom) of the same whole-hemisphere Visium v2 section (APP-KI). Twelve clusters were annotated to anatomical regions using marker gene expression and the Allen Mouse Brain Atlas. **(b)** Representative spatial maps of disease-associated astrocyte (DAA), disease-associated microglia (DAM), glycolysis (GO:0006096), oxidative phosphorylation (GO:0006119) and tricarboxylic acid (TCA) cycle (GO:0006099) module scores in WT and APP-KI hemisphere sections. Colour scales below the maps are shared between genotypes for each score and truncated at the 2nd and 98th percentiles of all spots. Signatures were defined by Habib et al.^62^ and Keren-Shaul et al.^61^, respectively, as previously applied to spatial transcriptomic data by Choi et al.^63^. **(c)** Regional genotype differences in module scores (n = 4 mice per group). Colours indicate the APP-KI − WT difference in each region estimated by a linear mixed model with animal as a random effect, divided by the standard deviation of the score across all spots. Benjamini– Hochberg correction was applied across the 12 regions; a false discovery rate (FDR) < 0.05 was considered significant. Asterisks indicate *FDR < 0.05, **FDR < 0.01 and ***FDR < 0.001. Spot-level distributions are shown in Supplementary Fig. 6. Statistics for all regions and scores are provided in Supplementary Table 11c. **(d)** Representative H&E-stained section (left) and spatial cell-type cluster map (right) of the hippocampus from Visium HD profiling. **(e)** Uniform manifold approximation and projection (UMAP) embedding of hippocampal Visium HD bins, coloured by annotated cell-type cluster. Ten clusters comprised two astrocyte populations, CA3 and CA1 pyramidal neurons, dentate gyrus granule neurons, GABAergic interneurons, microglia, oligodendrocytes and two unannotated clusters. **(f)** UMAP embedding as in e, coloured by genotype (APP-KI, orange; WT, blue). **(g)** Heatmap of scaled mean expression of selected marker genes used to annotate hippocampal cell-type clusters. Rows represent marker genes and columns represent clusters. The left annotation indicates the cell-type categories associated with the markers. **(h)** Gene set enrichment analysis (GSEA) of pseudobulk differential expression between APP-KI (*n* = 3) and WT (*n* = 2) mice within each annotated hippocampal cluster. Genes were ranked by the DESeq2 Wald statistic. The heatmap shows normalized enrichment scores (NES) for selected Gene Ontology Biological Process terms; positive and negative values indicate enrichment among genes with higher and lower expression in APP-KI mice, respectively. Colours indicate NES capped at ±3 for terms with FDR < 0.10; grey indicates FDR ≥ 0.10 or insufficient gene-set size. Symbols indicate †0.05 ≤ FDR < 0.10, *FDR < 0.05, **FDR < 0.01 and ***FDR < 0.001. The two unannotated clusters were excluded from this analysis. Abbreviations: APP-KI, *App*^NL-G-F^ knock-in; WT, wild type; TH, thalamus; WM, white matter; CTXsp+OLF, cortical subplate and olfactory areas; HPC, hippocampus; HY, hypothalamus; CP, caudoputamen; sAMY, striatum-like amygdalar nuclei; EPI, epithalamus; LV, lateral ventricle; CA1/CA3, cornu ammonis 1/3; DG, dentate gyrus; GABA-IN, GABAergic interneurons; HD, high definition.

We next profiled the hippocampus at higher spatial resolution using Visium HD (high definition; APP-KI, *n* = 3; WT, *n* = 2). Spatial mapping, uniform manifold approximation and projection (UMAP) visualization and marker expression identified eight annotated cell-type clusters, including two astrocyte-enriched clusters, microglia- and oligodendrocyte-enriched clusters and four neuron-enriched clusters, together with two unannotated clusters (Fig. 5d–g). Pseudobulk pathway analysis of the eight annotated clusters showed significant enrichment of complement activation and neuroinflammatory response signalling in five and seven clusters, respectively (FDR < 0.05; Fig. 5h). Reduced expression of mitochondrial pathway genes was detected in astrocyte-, microglia- and oligodendrocyte-enriched spatial clusters (Fig. 5h; Supplementary Table 11a,b). Complex I assembly showed an exploratory negative enrichment in γ-aminobutyric acid (GABA)-releasing interneurons (FDR = 0.075), whereas cornu ammonis 3 (CA3) and dentate gyrus neuronal clusters showed no significant mitochondrial pathway changes (Supplementary Table 11b). These findings identify reduced mitochondrial pathway signatures in glia-enriched hippocampal clusters, providing a cellular context for the MRS-detected metabolic alterations.

### Hyperpolarized MRS detects reduced bicarbonate labelling in APP-KI mice

Hyperpolarized [1-^13^C]pyruvate MRS in 6-month-old APP-KI mice showed a lower bicarbonate-to-pyruvate signal ratio than in WT mice (*P* = 0.043), while the lactate-to-pyruvate ratio did not differ significantly (*P* = 0.237). The bicarbonate-to-lactate ratio was also reduced (*P* = 0.010; Fig. 6). These findings are consistent with reduced PDH-linked oxidative pyruvate metabolism. Notably, a previous study reported reduced hyperpolarized lactate and bicarbonate signals following pharmacological inhibition of MCT1/4 in healthy mice, suggesting that alterations in monocarboxylate transport can influence these metabolic readouts^21^.

**Fig. 6.**
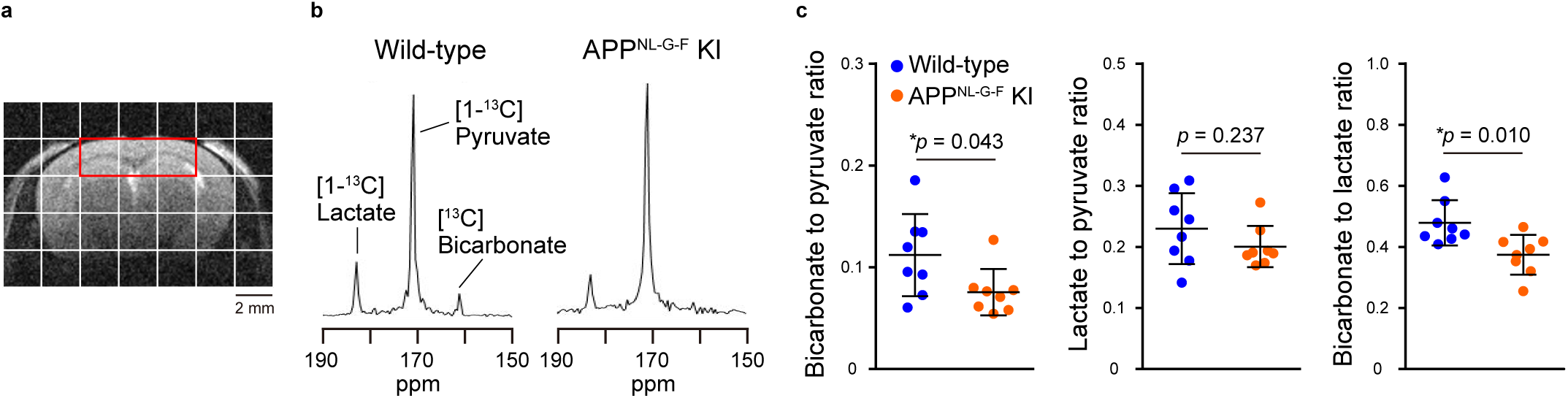
Hyperpolarized [1-^13^C]pyruvate MRS in APP-KI mice. Comparison of pyruvate metabolism in hippocampus-including regions between wild-type and APP-KI mice, evaluated by MRSI using HP [1-^13^C]pyruvate. (**a**) Coronal T_2_-weighted MR image of mouse brain, with red squares indicating VOI including the hippocampus and cortex. (**b**) Summed ^13^C spectra from 0 to 60 s after acquisition start, extracted from hippocampus and cortex-including VOI of 6-month-old awake APP-KI (right) and wild-type (left) mice administered HP [1-^13^C]pyruvate, acquired using an FID-based ^13^C MRSI sequence. (**c**) Quantification of the [^13^C]bicarbonate-to-[1-^13^C]pyruvate ratio (left), [1- ^13^C]lactate-to-[1-^13^C]pyruvate ratio (middle), and [^13^C]bicarbonate-to-[1-^13^C]lactate ratio (right) in hippocampus and cortex-including regions of 6-month-old awake APP-KI and wild-type mice, determined from peak heights measured in spectra summed over the acquisition period. Lines represent mean ± standard deviation; n = 8 mice per group. Statistical comparisons were performed using unpaired *t*-tests. Asterisks indicate \**P* < 0.05. Abbreviations: APP-KI, *App*^NL-G-F^ knock-in; MRS, magnetic resonance spectroscopy; MRSI, magnetic resonance spectroscopic imaging; HP, hyperpolarized; VOI, volume of interest; FID, free induction decay.

### Lower LC neuronal counts are associated with glial and mitochondrial protein changes in the PCC

Finally, given the role of noradrenergic signalling in glial and metabolic regulation, we examined whether LC neuronal integrity was associated with cortical glial and metabolic signatures in human postmortem tissue. We quantified tyrosine hydroxylase (TH)-positive LC neurons in cases with matched PCC proteomics (AD, *n* = 7; progressive supranuclear palsy (PSP), *n* = 6; control, *n* = 6), including PSP as a disease comparator with LC degeneration^22^. LC counts were lower in AD (median 28, range 9–114; *P* = 0.0012) and PSP (median 87.5, range 21–194; *P* = 0.041) than in controls (median 192.5, range 146–298; Fig. 7a).

**Fig. 7.**
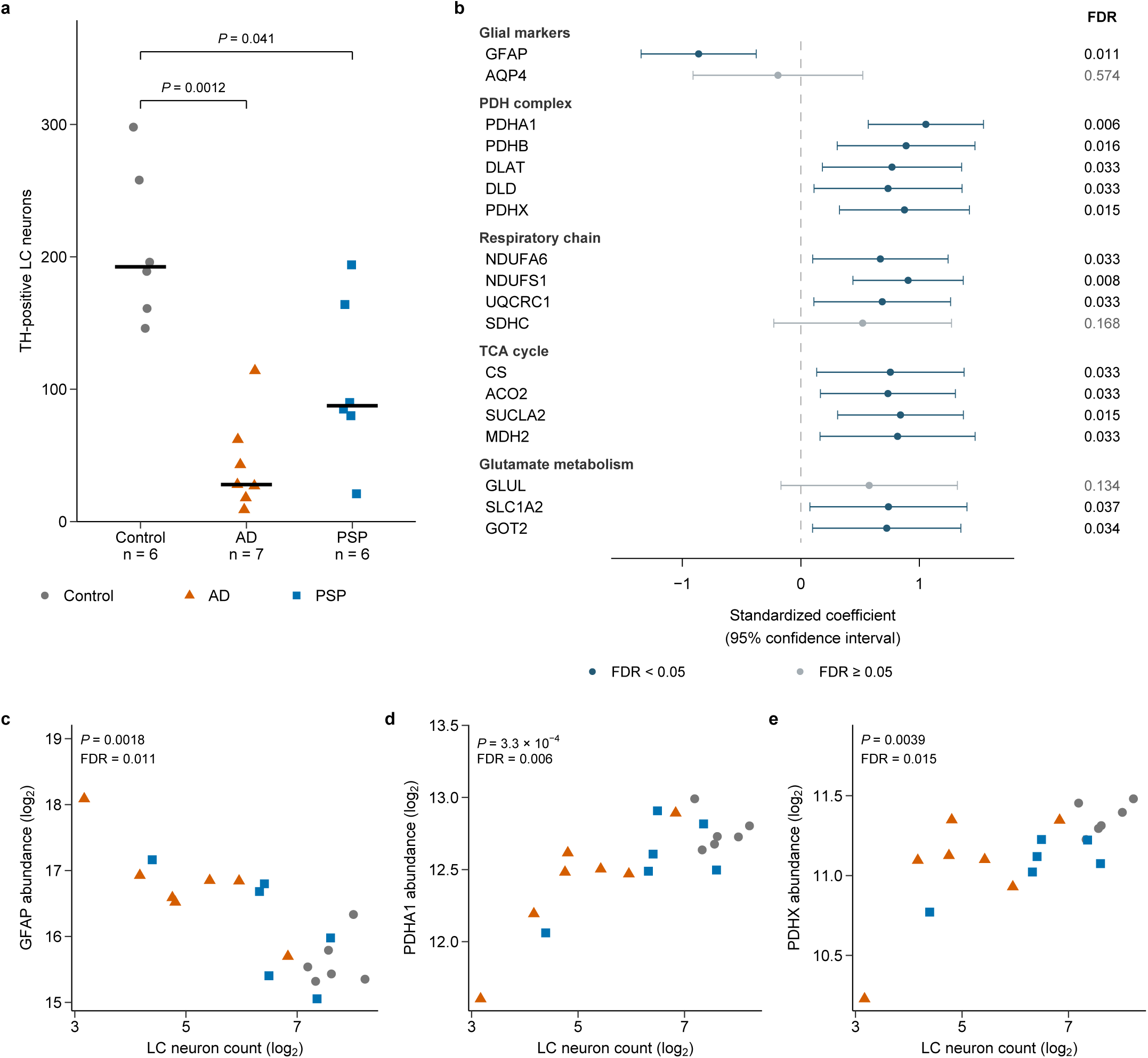
LC neuronal counts are associated with glial and mitochondrial metabolic proteins in the PCC. (**a**) TH-positive LC neuron counts, summed across the left and right sides, in control (*n* = 6), AD (*n* = 7) and PSP (*n* = 6) autopsy cases with matched PCC proteomics. Points represent individual cases; horizontal lines indicate medians. *P* values are from two-sided exact Mann– Whitney tests against controls and are unadjusted for the two group comparisons. (**b**) Associations between LC neuronal counts and the abundance of 18 selected PCC proteins, grouped by biological function. Points and horizontal bars show standardized regression coefficients and pointwise 95% confidence intervals from ordinary least-squares models of z(log_2_ protein abundance) as a function of z(log_2_ LC count), adjusted for diagnosis as a categorical covariate (control, AD or PSP). Standardization uses the mean and sample standard deviation across the 19 cases. Models include diagnosis-specific intercepts and a common LC coefficient; positive coefficients indicate higher protein abundance with higher LC counts after diagnosis adjustment. Two-sided *P* values for LC coefficients were BH-corrected across the 18 protein tests; dark blue, FDR < 0.05; grey, FDR ≥ 0.05. All 18 proteins were available in all 19 cases. Functional grouping does not imply cell-type specificity. (**c–e**) Individual-protein scatter plots for three proteins included in b: PCC GFAP (**c**), PDHA1 (**d**) and PDHX (**e**) abundance plotted against LC neuronal counts. Both axes are log_2_-transformed. Grey circles, controls; orange triangles, AD; blue squares, PSP. Points show observed values; displayed P and FDR values are from the diagnosis-adjusted models in b. All three panels show the same 19 cases used in b; the six control cases contribute once to the combined analysis. The 18-protein analysis is exploratory. Abbreviations: LC, locus coeruleus; PCC, posterior cingulate cortex; TH, tyrosine hydroxylase; AD, Alzheimer’s disease; PSP, progressive supranuclear palsy; FDR, false discovery rate; BH, Benjamini–Hochberg; GFAP, glial fibrillary acidic protein.

Across the 19 PCC cases, diagnosis-adjusted ordinary least-squares regression identified associations between LC neuronal counts and 15 of 18 selected proteins (BH FDR < 0.05; Fig. 7b). Lower LC counts were associated with higher GFAP abundance and lower abundance of multiple mitochondrial proteins, including all five tested PDH-complex components, respiratory-chain subunits NDUFA6, NDUFS1 and UQCRC1, TCA-cycle enzymes CS, ACO2, SUCLA2 and MDH2, and the mitochondrial aminotransferase GOT2, as well as the astrocytic glutamate transporter SLC1A2. Representative standardized LC coefficients were −0.86 for GFAP (95% confidence interval, −1.35 to −0.38; FDR = 0.011), 1.06 for PDHA1 (0.57 to 1.54; FDR = 0.006) and 0.87 for PDHX (0.33 to 1.42; FDR = 0.015; Fig. 7c–e). These estimates describe the LC–protein association after accounting for mean differences between diagnostic groups. Together, the PCC findings associate lower LC neuronal counts with greater GFAP abundance and reduced abundance of mitochondrial metabolic machinery.

## Discussion

Our earlier finding of elevated lactate and *myo*-inositol in the human AD PCC prompted investigation of the molecular and cellular changes accompanying this phenotype^11^. Here, human PCC proteomics and independent RNA-seq converged on reduced mitochondrial oxidative-metabolism signatures. The APP-KI hippocampus shared both mitochondrial proteomic signatures and MRS-detected lactate and myo-inositol alterations with the human AD PCC. This concordance supports the relevance of the APP-KI hippocampus for further investigation of AD-associated metabolic dysfunction. Hyperpolarized MRS provided in vivo evidence of reduced oxidative pyruvate metabolism, while **s**patial transcriptomics and proteomic network analysis implicated glial cells, particularly astrocytes, in the associated metabolic phenotype.

The PCC findings extend earlier evidence of reduced cytochrome oxidase activity^23^, lower respiratory-chain transcripts in neurons and proteins in tissue^24^, and altered mitochondrial transcripts in astrocytes^25^. By performing pathway analyses of independent PCC proteomic and transcriptomic datasets, we identified coordinated reductions across oxidative phosphorylation, respiratory-chain assembly and the TCA cycle. In ROSMAP PCC RNA-seq, mitochondrial scores tracked neuropathological severity, and TCA-cycle scores were associated with antemortem MMSE within clinical AD. Together, these findings reveal a coordinated reduction in molecular signatures of mitochondrial oxidative metabolism at both the transcript and protein levels in the AD PCC.

Beyond the molecular changes in human postmortem tissue, hyperpolarized [1-^13^C]pyruvate MRS provided in vivo evidence of altered oxidative pyruvate metabolism in APP-KI mice. At 6 months, APP-KI mice showed lower bicarbonate-to-pyruvate and bicarbonate-to-lactate signal ratios without a significant change in the lactate-to-pyruvate signal ratio, consistent with reduced PDH-linked conversion of pyruvate to bicarbonate. This finding parallels the negative enrichment of the PDH-mediated acetyl-CoA biosynthesis from pyruvate pathway in human PCC proteomics and independent RNA-seq data. These measurements provide information on pyruvate metabolism that cannot be inferred from lactate concentrations measured by conventional ^1^H-MRS alone. Previous hyperpolarized MRS studies reported increased pyruvate-to-lactate labelling in 3-month-old APP-KI mice^26^ and hAPP-J20 mice^27^, contrasting with the reduced bicarbonate labelling observed at 6 months in our study. A previous study using the same APP-KI model also reported early mitochondrial hypermetabolism followed by reduced hippocampal oxidative phosphorylation-related transcriptional signatures from 6 months of age^5^. Notably, hippocampal lactate concentrations in our study were elevated at 3, 10 and 13 months. Because lactate levels reflect the balance between production, utilization and transport, lactate elevation observed at multiple ages may accompany different underlying metabolic alterations across disease stages.

The regional and cellular context of this metabolic phenotype was examined through spatial profiling. Visium v2 revealed widespread increases in glycolysis-related transcriptional signatures and decreases in mitochondrial pathway signatures across multiple brain regions, including the hippocampus. Higher-resolution spatial profiling further identified reduced mitochondrial pathway signatures in astrocyte-, microglia- and oligodendrocyte-enriched spatial clusters in the APP-KI hippocampus. Although reduced mitochondrial gene expression has been reported in human AD PCC neurons^24^, our findings highlight glial involvement in the APP-KI hippocampus.

Human proteomic network analysis further supports an astrocyte-associated context for the oxidative metabolic changes. The downregulated M4 module was enriched for astrocyte markers, with mitochondrial proteins covarying with ATP-dependent glutamine synthetase and synaptic components. Although RNA-seq also showed negative enrichment of oxidative metabolism-related pathways, no corresponding co-expression module was identified. Most oxidative phosphorylation genes were excluded by CEMiTool’s variance-based filter, potentially limiting the detection of such a module. The difference between the proteomic and transcriptomic networks may also reflect tighter coordination of protein-complex abundance at the protein level^28^. The organization of M4 is consistent with human ^13^C-MRS evidence for astrocytic oxidative metabolism and the contribution of glutamate–glutamine cycling to neuronal glutamate replenishment^29^. In APP-KI mice, hippocampal glutamine increased over the same age range as myo-inositol, and because glutamine synthetase is largely astrocytic, this concurrent rise is consistent with an astrocytic contribution to the metabolite changes detected in vivo. Glutamate uptake is also linked to astrocytic glycolysis and lactate production, which underlie the proposed astrocyte–neuron lactate shuttle^30^. Consistent with this, hyperpolarized [1-^13^C]pyruvate conversion is sensitive to astrocyte-associated lactate metabolism, as pharmacological blockade of astrocytic monocarboxylate transporters reduces pyruvate-to-lactate and pyruvate-to-bicarbonate labelling in the mouse brain in vivo^21^. Previous large-scale AD brain proteomics identified an upregulated astrocyte- and microglia-enriched module involving glucose metabolism, including glycolysis, that was associated with AD pathology and cognitive impairment^4^. Our findings extend these observations by identifying a downregulated astrocyte-marker-enriched module linking mitochondrial oxidative metabolism with neurotransmitter handling.

Beyond local metabolic remodelling, our findings associate LC neuronal counts with glial and mitochondrial protein abundance in the PCC. LC degeneration accompanies AD pathological progression^31^, while noradrenergic signalling regulates astrocytic lactate metabolism and microglial responses to amyloid-β (Aβ)^32,33^. Across AD, PSP and controls, lower LC neuronal counts were associated with higher PCC GFAP abundance and lower abundance of PDH-complex, respiratory-chain and TCA-cycle proteins after adjustment for diagnostic group. The association with PDH-complex proteins is consistent with the PDH-linked metabolic alteration suggested by hyperpolarized MRS in APP-KI mice. In a related clinical study, Prokopiou et al.^34^ linked lower LC activity and functional connectivity to higher plasma tau phosphorylated at threonine 217 (p-tau217), with these associations becoming more pronounced as GFAP increased. These LC–PCC associations raise the possibility that altered noradrenergic input contributes to cortical metabolic changes, although this mechanism remains to be tested.

Several limitations warrant consideration. Human postmortem analyses are cross-sectional, limiting causal and temporal interpretations. Proteomic concordance between human PCC and mouse hippocampus supports shared metabolic alterations, although regional and age-related differences limit direct comparisons. Finally, the LC associations require independent validation with more comprehensive neuronal quantification.

Together, these findings identify convergent alterations in mitochondrial energy metabolism across human AD and APP-KI mice, alongside in vivo evidence of reduced oxidative pyruvate metabolism in APP-KI mice. These results highlight the potential involvement of glial cells, particularly astrocytes, in AD-associated metabolic dysfunction.

## Online Methods

### Postmortem human brain tissue

Autopsied brain tissue was obtained from the Brain Bank of the Niigata University Brain Research Institute. All autopsies were performed with written informed consent from the legal next of kin or from an individual holding power of attorney, and the study using autopsy material was approved by the National Institutes for Quantum Science and Technology (QST) Certified Review Board (approval number N22-006). We included seven AD cases that histologically exhibited abundant amyloid-β (Aβ)-positive senile plaques and phosphorylated tau-positive neurofibrillary tangles and neuropil threads in the cerebral neocortex, consistent with Braak stage IV/C or higher^35,36^. The clinicopathological characteristics of the patients with AD, including age, sex, disease duration and neuropathological assessment according to the National Institute on Aging–Alzheimer’s Association (NIA-AA) criteria^37^, are summarized in Supplementary Table 12. We also included nine control cases without neurological disorders and eight patients with PSP. All control and PSP cases exhibited Braak stage III/C or lower. Details of the control and PSP cases and their PCC proteomic data have been described previously^22^. No phosphorylated TAR DNA-binding protein of 43 kDa (TDP-43) was detected in the AD, control or PSP cases, except for one AD case showing TDP-43-positive inclusions in the hippocampus, consistent with stage 2 limbic-predominant age-related TDP-43 encephalopathy neuropathologic change (LATE-NC)^38^. Phosphorylated α-synuclein was not evident in any group, except in two PSP cases showing a small number of Lewy bodies and neurites limited to the medulla oblongata without overt neuronal loss in the dorsal motor nucleus of the vagal nerve, considered subclinical. None of the AD, PSP or control cases exhibited severe ischaemic or haemorrhagic brain injury, hypoxic encephalopathy, or central nervous system (CNS) infectious or metabolic disorders.

At autopsy, the right cerebral hemisphere was sliced coronally (perpendicular to the orbitomeatal line), snap-frozen and stored at −80 °C. The left cerebral hemisphere, cerebellum and brainstem were immersed in 10% buffered formalin. Frozen tissue samples of the PCC were collected from the frozen cingulate cortices at coronal levels approximately corresponding to the splenium of the corpus callosum and were used for SWATH-MS-based quantitative proteomics (AD, *n* = 7; control, *n* = 9). Immunohistochemistry was performed on formalin-fixed, paraffin-embedded (FFPE) sections of the left PCC at the corresponding coronal levels, as well as on sections of the LC obtained from horizontally sectioned rostral pons tissue.

### ROSMAP bulk RNA-seq cohort and diagnostic frameworks

Bulk RNA-seq data from the posterior cingulate cortex (PCC) were obtained from 450 ROSMAP participants^20^. Tissue samples and clinical and neuropathological data were provided by the Rush Alzheimer’s Disease Center, Rush University Medical Center, Chicago, and the RNA-seq data were obtained from the AD Knowledge Portal. Two complementary diagnostic frameworks were applied.

For the clinical framework (cogdx_group), diagnoses were assigned at death by study physicians on the basis of clinical history and detailed neuropsychological evaluation while blinded to neuropathological results^39^. Clinical AD dementia was defined as probable or possible AD dementia (cogdx = 4 or 5), based on the National Institute of Neurological and Communicative Disorders and Stroke–Alzheimer’s Disease and Related Disorders Association (NINCDS–ADRDA) criteria^40^ (*n* = 165), and cognitively normal controls (no cognitive impairment; NCI) were defined as cogdx 1 (*n* = 162), regardless of neuropathological findings. For the pathological framework (patho_dx), AD required cogdx 4 or 5 combined with a high (Braak ≥ V and CERAD = 3) or intermediate (Braak III–IV and CERAD ≥ 2) National Institute on Aging (NIA)–Reagan likelihood of AD^41^ (*n* = 117); NCI controls required cogdx 1 together with an absence of neuritic plaques (CERAD = 0) (*n* = 69).

### Mice

All animal procedures were performed in accordance with the National Research Council Guide for the Care and Use of Laboratory Animals and are reported in accordance with the Animal Research: Reporting of In Vivo Experiments (ARRIVE) guidelines 2.0^42^. Protocols were approved by the Animal Ethics Committee of the National Institutes for Quantum Science and Technology (approval number 22-1012-4). Fifty adult *App^NL-G-F^* knock-in AD model mice (3–18 months)^43^, 24 adult WT littermates (6–18 months), 109 adult rTg4510 tauopathy model mice (3–8 months)^44^, and 58 adult non-transgenic (non-Tg) littermates (3–8 months) were used. Both male and female mice were used in all experiments. Mice were maintained on a 12-h light/dark cycle with ad libitum access to a standard diet and water. Group sizes for each experiment are given in the corresponding sections below, in the figure legends and in Supplementary Table 13.

### Mouse tissue collection

Proteomics. Eighteen-month-old APP-KI and WT mice were deeply anaesthetized and killed by cervical dislocation, and brains were removed. Hippocampal tissue was dissected, snap-frozen and stored at −80 °C.

Tissue collection for spatial transcriptomics is described below.

### SWATH-MS-based quantitative proteomics

Frozen tissue from the PCC of patients with AD and control individuals (AD, *n* = 7; control, *n* = 9) and from the hippocampus of 18-month-old APP-KI and WT mice (APP-KI, *n* = 6; WT, *n* = 6) was homogenized in denaturing buffer (7 M guanidine hydrochloride, 500 mM Tris– HCl pH 8.5, 10 mM ethylenediaminetetraacetic acid (EDTA)) at 8,500 rpm with 20 zirconia beads (1.0 mm diameter) using a Precellys Evolution homogenizer (Bertin Instruments). Whole-tissue lysates (50 μg protein per sample) were prepared and analysed by liquid chromatography–tandem mass spectrometry (LC–MS/MS) as described previously^45^.

### Immunohistochemistry

For the human PCC analysis, double immunofluorescence labelling for GFAP and NDUFA6 was performed on FFPE sections from three control and three AD cases included in the SWATH-MS proteomic cohort. Left-hemisphere PCC sections at corresponding coronal levels were used, whereas proteomics used right-hemisphere tissue. Deparaffinized sections were immunostained with a rat monoclonal antibody against GFAP (13-0300; Invitrogen; 1:200) and a rabbit polyclonal antibody against NDUFA6 (FNab05610; FineTest; 1:1,000). High- and lower-magnification images were used for qualitative assessment of the distribution of NDUFA6 immunoreactivity relative to GFAP-positive profiles; no quantitative comparison of staining between diagnoses was performed. Immunolabelling was examined with a BZ-X710 microscope (Keyence).

### Locus coeruleus neuronal quantification

Pontine FFPE sections containing the LC were immunostained with a mouse monoclonal antibody against tyrosine hydroxylase (TH-16; Sigma-Aldrich, T2928; 1:2,000). TH-positive neurons with a diameter ≥ 20 μm were counted bilaterally and summed, as described previously^22^. Cases in which the LC was recorded as unsuitable for counting were excluded (three control and two PSP cases with PCC proteomic data). Counting was performed blinded to diagnosis.

### In vivo ^1^H-MRS

Single-voxel ^1^H-MRS was performed in the hippocampus and cortex on a 7 T Bruker scanner (20 cm bore, BioSpec, Avance-III; Bruker BioSpin) equipped with a two-channel phased-array cryogenic surface coil for transmission and reception (cryoprobe for mouse brain; Bruker BioSpin). Mice were anaesthetized with 1.5 ± 0.3% (v/v) isoflurane, and a point-resolved spectroscopy (PRESS) sequence with a repetition time (TR) of 4,000 ms and an echo time (TE) of 20 ms was applied. Signals were acquired from volumes of interest placed in the dorsal hippocampus (1.2 × 2.0 × 2.2 mm^3^) and cortex (1.2 × 2.5 × 3.0 mm^3^) and averaged over 256 acquisitions. Water suppression was achieved with the variable power and optimized relaxation delays (VAPOR) module^46^.

Spectra were processed with LCModel (v6.3-1L, Stephen Provencher)^47^ without preprocessing and fitted to a linear combination of a simulated basis set of 21 metabolites: alanine, aspartate, phosphocholine, creatine, phosphocreatine, GABA, glutamine, glutamate, glutathione, glycine, *myo*-inositol, lactate, N-acetylaspartate (NAA), *scyllo*-inositol, taurine, glucose, N-acetylaspartylglutamate (NAAG), glycerophosphocholine, phosphorylethanolamine, serine and β-hydroxybutyrate. The unsuppressed water signal acquired from the same volume of interest was used as an internal reference for absolute quantification. The following metabolites were evaluated: lactate, *myo*-inositol, tNAA (NAA + NAAG), tCho (glycerophosphocholine + phosphocholine), aspartate, GABA, glucose, glutamate, glutamine, total creatine (tCr; creatine + phosphocreatine) and taurine.

### Awake-mouse preparation for MRS

For awake hyperpolarized [1-^13^C]pyruvate MRS, a custom-made headplate was affixed to the skull with dental adhesive and resin as described previously^48^, at least one week before the first imaging session. Mice were then acclimated to a custom-made magnetic resonance imaging (MRI)/MRS-compatible awake-mouse restraint device. Animals were anaesthetized with 1.5% (v/v) isoflurane only during insertion of the tail-vein catheter and positioning in the restraint device via the headplate, and the isoflurane supply was turned off at least 20 min before in vivo MRS acquisition.

### In vivo hyperpolarized ^13^C magnetic resonance spectroscopic imaging

[1-^13^C]Pyruvic acid (24 μl; CLM-8077-PK, Cambridge Isotope Laboratories) was mixed with 25 mM OX063 (Polarize) and hyperpolarized at 1.3 K and 6.7 T in a SpinAligner dissolution dynamic nuclear polarization (dDNP) polarizer (Polarize). Microwave irradiation was applied at 187.84 GHz and 22 mW, following optimization performed on the same sample and system^48^. At maximum polarization, dissolution was performed with 4 ml of a superheated alkaline buffer containing 40 mM tris(hydroxymethyl)aminomethane (819620, MP Biomedicals), 50 mM sodium chloride (191-01665, FUJIFILM Wako), 0.27 mM EDTA (195173, MP Biomedicals) and 80 mM sodium hydroxide (31511-05, Nacalai Tesque). The final pyruvate concentration after dissolution and neutralization was approximately 80 mM.

All hyperpolarized magnetic resonance (MR) studies were performed on a BioSpec 3 T preclinical scanner (Bruker BioSpin) with a ^1^H–^13^C dual-tuned surface transmit/receive coil (Takashima Seisakusho). The ^13^C coil was loop-shaped (18 × 22 mm) and bent along its 18 mm axis to conform to a cylindrical surface 24 mm in diameter; the ^1^H coil was butterfly-shaped (32 × 24 mm) and bent along its 32 mm axis to fit the same cylindrical surface.

After positioning inside the magnet, axial, sagittal and coronal two-dimensional localizer images were acquired with a rapid acquisition with relaxation enhancement (RARE) sequence (TR = 2,400 ms, TE = 12 ms, RARE factor = 10, four averages, one repetition, field of view 24 × 24 mm, matrix 192 × 192, flip angle 90°). A slab for magnetic resonance spectroscopic imaging (MRSI) was positioned across the coronal plane to cover the whole brain, or across the axial plane to primarily cover the hippocampus and cortex (Supplementary Fig. 7). Targeted regions were shimmed using Bruker MAPSHIM based on acquired B_0_ field maps, followed by localized first- and second-order shimming where necessary, reducing the localized proton linewidth to approximately 10–20 Hz (mean, approximately 15 Hz) within the targeted brain volume.

Each mouse was injected intravenously via a tail-vein catheter with hyperpolarized [1- ^13^C]pyruvate solution at a dose of 10 μl per g body weight and at a rate of approximately 50 μl s^−1^. ^13^C MRSI was performed with a free induction decay (FID)-based chemical shift imaging sequence to map the time course and spatial distribution of metabolites, starting 4 s after injection of hyperpolarized [1-^13^C]pyruvate (flip angle 10°, TR 85 ms, TE 1.245 ms, coronal slice thickness 8 mm, five temporal repetitions, field of view 24 × 24 mm, matrix 12 × 12; total scan time 61 s). Hyperpolarized ^13^C metabolite maps were generated by extracting peak heights for each metabolite and overlaying them on proton T_2_-weighted images. For quantitative analysis, spectra were obtained by summing all five temporal repetitions. MRSI data were analysed in magnitude mode with custom programs written in MATLAB (vR2020b): FID data were Fourier transformed, spectra from regions of interest were extracted and, where multiple regions or temporal repetitions were involved, summed; metabolite ratios were then quantified from peak heights.

### Spatial transcriptomics: tissue processing and library preparation

Eight mice were used: APP-KI (*n* = 4) and age-matched WT (*n* = 4). All animals underwent ^1^H-MRS and hyperpolarized ^13^C MRSI at 6 months of age, followed by tissue collection for spatial transcriptomics at 7 months of age. This design enabled comparison of in vivo metabolic imaging and subsequent spatial transcriptomic profiles within the same animals. Mice were deeply anaesthetized and perfused with PBS followed by 4% PFA using a perfusion fixation toolkit (INT-PFTS-M, GenoStaff). Brains were dissected, postfixed in 4% PFA at room temperature for up to one week, washed in PBS and stored in 70% ethanol at 4 °C. Fixed samples were embedded in paraffin and sectioned at 10 μm on a microtome.

From each animal, one hemisphere section was used for Visium v2 profiling (55-μm-diameter spots, 100-μm centre-to-centre spacing; eight sections in total; APP-KI *n* = 4, WT *n* = 4). From a subset of these animals (APP-KI *n* = 3, WT *n* = 3), an additional hippocampus-containing section was processed for Visium HD profiling (2-μm-resolution barcoded grid, binned to 8 μm for downstream analysis). One WT section was excluded at quality control, leaving APP-KI *n* = 3 and WT *n* = 2 for Visium HD analysis.

For Visium v2, FFPE sections were deparaffinized and stained with haematoxylin and eosin (H&E) according to the manufacturer’s protocol (CG000520, Rev B), and images were acquired using an APX100 microscope (Evident). Libraries were prepared using Visium CytAssist Spatial Gene Expression reagents with the Visium Mouse Transcriptome Probe Set v1.0 and the Dual Index Kit TS Set A (10x Genomics), following the manufacturer’s user guide (CG000495, Rev E). Library quality was assessed using an Agilent 2200 TapeStation, and DNA concentrations were measured using a Qubit 4 Fluorometer with the Qubit 1X dsDNA HS Assay Kit (Thermo Fisher Scientific). Libraries were sequenced on a DNBSEQ-G400 platform (MGI), with 28-bp read 1 and 100-bp read 2.

For Visium HD, sections used for spatial profiling were deparaffinized and H&E-stained following the Visium HD FFPE Tissue Preparation Handbook (CG000684, Rev A). Images were acquired using a VS200 slide scanner (Evident), following the Visium HD Spatial Applications Image Guidelines (CG000688, Rev A). Libraries were prepared using the Visium HD Mouse Transcriptome, 6.5 mm kit and the Dual Index Kit TS Set A (10x Genomics), following the manufacturer’s user guide (CG000685, Rev A). Library quality was assessed using an Agilent 4200 TapeStation, and DNA concentrations were measured using a Qubit 4 Fluorometer with the Qubit 1X dsDNA HS Assay Kit. Libraries were sequenced on a NovaSeq X Plus platform (Illumina), with paired-end 150-bp reads.

Raw FASTQ files and corresponding H&E images from both platforms were processed using Space Ranger (v3.0.1; 10x Genomics).

### Data analysis

#### Proteomic data processing and differential abundance analysis

Differential protein abundance (AD versus control; APP-KI versus WT) was assessed by two-sided Student’s *t*-test on protein intensities, with log_2_ fold changes calculated from group means and *P* values adjusted across proteins by the BH method^49^. Mouse proteins quantified in at least three samples per group were analysed. No human protein reached adjusted *P* < 0.05; proteins with nominal *P* < 0.05 and |log_2_FC| > 0.1 are reported as exploratory (Fig. 1b), and pathway-level inference relied on GSEA and GSVA.

#### ROSMAP bulk RNA-seq processing and differential expression analysis

Raw counts were analysed with DESeq2 (v1.46.0)^50^. After exclusion of samples with missing covariates and one sample from a single-sample sequencing batch, 450 samples remained. Protein-coding genes with at least 10 counts in at least 10 samples were analysed. Differential expression between pathologically defined AD and NCI cases was tested, adjusting for age at death, postmortem interval (PMI), RIN, sequencing batch and sex; differentially expressed genes were defined by FDR < 0.05 and |log_2_FC| > 0.1.

#### Gene set enrichment and over-representation analyses

GSEA^51^ of GO BP terms was performed with gseGO in clusterProfiler (v4.14.6)^52^, ranking all quantified proteins by the *t* statistic or genes by the DESeq2 Wald statistic (gene sets of 10– 500 genes; nPermSimple = 10,000; seed = 42). Terms with FDR < 0.05 were considered significant; representative terms are displayed, and full results are provided in the Supplementary Tables. Co-expression modules were annotated with Molecular Signatures Database (MSigDB) gene sets^53^ (see below).

#### Gene set variation analysis

GSVA (v2.0.7)^54^ scores (Gaussian kernel) were calculated for four GO BP gene sets selected a priori: oxidative phosphorylation (GO:0006119), TCA cycle (GO:0006099), mitochondrial respiratory chain complex assembly (GO:0033108) and glycolytic process (GO:0006096). Differences between AD and controls were tested with limma (v3.62.2)^55^ and BH correction.

For proteomics, scores were computed from log_2_-transformed, median-normalized abundances after removal of proteins missing in more than 50% of samples and k-nearest-neighbour imputation, and diagnosis was the only predictor. For RNA-seq, GSVA scores were computed from variance-stabilized expression across all 450 samples, and limma models included age at death, PMI, RIN, sequencing batch and sex. Group comparisons and neuropathological association analyses used the 186 pathologically defined AD and NCI cases, whereas MMSE associations were assessed within the 165 clinically diagnosed AD cases. Covariate-adjusted scores in Fig. 2c are shown for visualization only.

#### Weighted gene co-expression network analysis

Co-expression modules were identified with CEMiTool (v1.30.0)^56^, which implements weighted gene co-expression network analysis^57^, using default settings. Inputs were log_2_-transformed protein abundances (*n* = 16) and variance-stabilized RNA-seq counts of the 186 pathologically defined samples after removal of ribosomal protein genes and regression of sequencing batch, RIN, age at death, PMI and sex with limma::removeBatchEffect. The variance filter retained 510 proteins and 1,848 genes, and soft-thresholding powers were selected automatically (β = 7 and 4), yielding five protein and 11 gene modules. Modules were annotated by over-representation analysis against MSigDB (v7.4) GO BP gene sets and by one-sided Fisher’s exact tests for curated cell-type marker genes^4^, with BH correction within each cell type. Differential module activity between AD and controls was assessed with CEMiTool’s built-in GSEA, and RNA-seq module eigengenes were regressed on Braak stage, CERAD score or MMSE, adjusting for age at death, PMI, RIN and sex, with BH correction within each trait.

#### Association of pathway enrichment scores with cognition and neuropathology

Within clinical AD (*n* = 165), MMSE scores at the last evaluation were regressed on GSVA scores for the TCA cycle and oxidative phosphorylation, adjusting for age at death, PMI, RIN, sequencing batch and sex, with BH correction across the two pathways.

Associations with CERAD score (recoded so that higher values indicate more neuritic plaques) and Braak stage were tested in the 186 pathologically defined samples by proportional-odds ordinal logistic regression (MASS::polr) with the same covariates, with BH correction within each trait.

#### Cross-species comparison of human and mouse proteomes

Human and mouse proteomes were compared across 4,881 one-to-one orthologs (babelgene v22.9) quantified in both species. Genome-wide concordance was assessed by rank–rank hypergeometric overlap (RRHO2 v1.0)^58^ of orthologs ranked by signed −log_10_ *P*. For the 700 orthologs meeting the exploratory human threshold, directional agreement was tested with a two-sided binomial test, and log_2_ fold changes were compared by Spearman correlation. Pathway-level concordance was assessed by Spearman correlation of GSEA NES across the 3,717 GO BP terms tested in both species.

#### Visium v2 spatial transcriptomic analysis

Space Ranger output was processed with Seurat (v5.0.2)^59^ using SCTransform normalization^60^. Unsupervised clustering identified 12 clusters, which were annotated to anatomical regions by cross-referencing cluster-specific marker gene expression with the Allen Mouse Brain Atlas: cortex 1 (Cortex 1), thalamus, white matter (WM), cortical subplate and olfactory areas (CTXsp+OLF), hippocampus (HPC), hypothalamus (HY), cortex 2 (Cortex 2), meninges, caudoputamen (CP), striatum-like amygdalar nuclei (sAMY), epithalamus (EPI) and lateral ventricle (LV).

Module scores were calculated per spot with AddModuleScore for three energy metabolism-related GO BP gene sets (GO:0006119, oxidative phosphorylation; GO:0006099, TCA cycle; GO:0006096, glycolysis) and for DAM (*Lpl*, *Cst7*, *Axl*, *Itgax*, *Spp1*, *Cd9*, *Ccl6*, *Csf1*)^61^ and DAA (*Ggta1*, *Gsn*, *Osmr*, *Vim*, *Serpina3n*, *Ctsb*, *Gfap*)^62^ signatures, as applied to spatially resolved transcriptomic data previously^63^.

Because spots from the same animal are not statistically independent, module scores were modelled with a linear mixed model including a genotype (APP-KI versus WT) × region interaction and animal as a random effect (lme4/lmerTest). Pairwise contrasts between APP-KI and WT within each region were extracted with emmeans using asymptotic (z) degrees of freedom, with BH correction applied across the 12 regions; contrasts were considered significant at FDR < 0.05 (Supplementary Table 11c). For display (Fig. 5c), each contrast was divided by the standard deviation of the corresponding score across all spots.

#### Visium HD spatial transcriptomic analysis

For Visium HD analyses, 8-μm bins corresponding to the hippocampus were extracted and normalized using SCTransform (Seurat v5.0.2), with the percentage of mitochondrial transcripts included as a regression variable. Principal component embeddings were corrected for interindividual variation using Harmony (v1.2.0)^64^. Seurat graph-based clustering yielded 10 clusters, which were annotated on the basis of known marker genes as two astrocyte populations, cornu ammonis 1 (CA1) and CA3 pyramidal neurons, dentate gyrus granule neurons, microglia, oligodendrocytes, GABAergic interneurons and two unannotated clusters. For each annotated cluster, bin-level counts were aggregated by biological sample to generate pseudobulk count matrices. Differential expression between APP-KI and WT was analysed using DESeq2 with a Wald test. For pathway analysis, genes were ranked according to the DESeq2 Wald statistic, and GSEA was performed separately within each cluster using GO BP gene sets.

#### Locus coeruleus neuronal counts and cortical protein associations

An exploratory panel of 18 proteins related to the glial and energy-metabolism findings was selected (GFAP, AQP4, PDHA1, PDHB, DLAT, DLD, PDHX, NDUFA6, NDUFS1, UQCRC1, SDHC, CS, ACO2, SUCLA2, MDH2, GLUL, SLC1A2 and GOT2); all were quantified in the 19 PCC cases with LC counts (AD, *n* = 7; PSP, *n* = 6; control, *n* = 6). LC counts in each disease group were compared with controls by two-sided exact Mann–Whitney tests. For each protein, log_2_ abundance was regressed on log_2_ LC count with diagnosis as a covariate (ordinary least squares, continuous variables standardized), and *P* values for the LC coefficient were adjusted across the 18 proteins by the BH method. PCC proteomic data and TH-positive LC neuron counts from PSP and control cases were previously reported^22^ (PRIDE accession PXD066064). The associations between LC neuron counts and cortical protein abundance are reported here for the first time.

#### Statistics

Statistical analyses of in vivo imaging data were performed in GraphPad Prism v6.0 (GraphPad Software). Comparisons between two groups used unpaired two-sided *t*-tests. For comparisons among three or more groups, one-way analysis of variance (ANOVA) followed by Dunnett’s multiple comparisons test was used, with wild-type mice at 6 and 12 months pooled as a single control group and each APP-KI age group compared against it. Correlations between metabolite concentrations were assessed with Pearson’s correlation coefficient. Group sizes (n) for the analyses shown in Fig. 4c, Fig. 6c and Supplementary Figs. 3, 4b and 5c are given in Supplementary Table 13.

All omics analyses were performed in R (v4.4.2). Key packages were DESeq2 (v1.46.0), clusterProfiler (v4.14.6), GSVA (v2.0.7), CEMiTool (v1.30.0), WGCNA (v1.74), limma (v3.62.2), MASS (v7.3.65), Seurat (v5.0.2), Harmony (v1.2.0), lme4/lmerTest, emmeans, org.Hs.eg.db (v3.20.0) and org.Mm.eg.db (v3.20.0).

Unless otherwise stated, statistical tests were two-sided and significance was defined as Benjamini–Hochberg (BH)-adjusted *P* value (false discovery rate, FDR) < 0.05. For pseudobulk GSEA of Visium HD clusters, statistical significance was defined as FDR < 0.05. The heatmap additionally displays exploratory findings with 0.05 ≤ FDR < 0.10; these are coloured and marked with a dagger (†), whereas asterisks indicate FDR < 0.05. All other GSEA results used FDR < 0.05.

## Data availability

The mass spectrometry raw data and DIA-NN-processed data have been deposited in the ProteomeXchange Consortium via the jPOST partner repository with the data set identifier PXD087601 (JPST005079) for human and PXD087402 (JPST005080) for mice^65^, and the raw and processed spatial transcriptomics data will be deposited in the Gene Expression Omnibus; both will be made publicly available upon publication.

ROSMAP posterior cingulate cortex bulk RNA-seq data and associated clinical and neuropathological metadata are available via the AD Knowledge Portal (https://adknowledgeportal.synapse.org/). The AD Knowledge Portal is a platform for accessing data, analyses, and tools generated by the Accelerating Medicines Partnership (AMP-AD) Target Discovery Program and other National Institute on Aging (NIA)-supported programs to enable open-science practices and accelerate translational learning. The data, analyses and tools are shared early in the research cycle without a publication embargo on secondary use. Data are available for general research use according to the following requirements for data access and data attribution (https://adknowledgeportal.synapse.org/Data%20Access). For access to the data resources described in this manuscript, see https://doi.org/10.7303/9618239.

## Code availability

Analysis code will be made available on GitHub upon publication.

## Acknowledgements

We thank the participants of the Religious Orders Study and the Rush Memory and Aging Project (ROSMAP) and the investigators and staff of the Rush Alzheimer’s Disease Center, Rush University Medical Center, Chicago, for their contributions to this research. The results published here are in whole or in part based on data obtained from the AD Knowledge Portal (NIH-NIA, U24AG061340). Study data were provided by the Rush Alzheimer’s Disease Center, Rush University Medical Center, Chicago. Data collection was supported through funding by NIA grants P30AG10161 (ROS), R01AG15819 (ROSMAP; genomics and RNAseq), R01AG17917 (MAP), R01AG36836 (RNAseq), P30AG072975 and the Illinois Department of Public Health (ROSMAP). Additional phenotypic data can be requested at www.radc.rush.edu.

This work was supported by AMED under grant number JP24dk0207063 (to Y.T.); the MEXT Promotion of Development of a Joint Usage/Research System Project: Coalition of Universities for Research Excellence Program (CURE) under grant number JPMXP1323015488, “Frontier of Spin Life Sciences (Spin-L)”; the JSPS Joint Research Program implemented in association with SNSF (JRPs) under grant number JPJSJRP22021510 (to Y.T.); JSPS KAKENHI grant numbers JP26K21969 and JP24K02411 (to Y.T.) and JP25K10974 (to M.O.); the Collaborative Research Project of the Brain Research Institute, Niigata University, under grant number 24014 (to M.O.); the AMED Strategic Research Program for Brain Sciences under grant number JP25wm0625126 (to A.K.); the AMED Moonshot Research and Development Program under grant number JP24zf0127012 (to A.K.); and the JSPS J-PEAKS Program under grant number JPJS00420240016.

## Author contributions

Y.T. designed the study. K.H. performed human proteomic and transcriptomic analyses and cross-species comparisons. M.O. conducted mouse experiments and MRS measurements. Y.M., Y.A. and Y.U. performed proteomic measurements. R.K., A.U., G.K. and H.E. performed mouse MRS analyses. K.BP. and T.R. performed bulk RNA-seq analyses. H.S. and A.K. performed neuropathological analyses and LC neuron counts. K.D., M.M. and J.N. contributed to the conception and interpretation of the study. Y.U. and Y.T. supervised the study. K.H. and M.O. wrote the original manuscript with contributions from all other authors. All authors reviewed and edited the manuscript.

## References

1. Scheltens, P. et al. Alzheimer’s disease. Lancet 397, 1577–1590 (2021).

2. Cunnane, S. C. et al. Brain energy rescue: an emerging therapeutic concept for neurodegenerative disorders of ageing. Nat. Rev. Drug Discov. 19, 609–633 (2020).

3. Salmon, E., Collette, F. & Bastin, C. Cerebral glucose metabolism in Alzheimer’s disease. Cortex 179, 50–61 (2024).

4. Johnson, E. C. B. et al. Large-scale proteomic analysis of Alzheimer’s disease brain and cerebrospinal fluid reveals early changes in energy metabolism associated with microglia and astrocyte activation. Nat. Med. 26, 769–780 (2020).

5. Naia, L. et al. Mitochondrial hypermetabolism precedes impaired autophagy and synaptic disorganization in App knock-in Alzheimer mouse models. Mol. Psychiatry 28, 3966– 3981 (2023).

6. Buckner, R. L. et al. Molecular, structural, and functional characterization of Alzheimer’s disease: evidence for a relationship between default activity, amyloid, and memory. J. Neurosci. 25, 7709–7717 (2005).

7. Reiman, E. M. et al. Preclinical evidence of Alzheimer’s disease in persons homozygous for the epsilon 4 allele for apolipoprotein E. N. Engl. J. Med. 334, 752–758 (1996).

8. Minoshima, S. et al. Metabolic reduction in the posterior cingulate cortex in very early Alzheimer’s disease. Ann. Neurol. 42, 85–94 (1997).

9. Gordon, B. A. et al. Spatial patterns of neuroimaging biomarker change in individuals from families with autosomal dominant Alzheimer’s disease: a longitudinal study. Lancet Neurol. 17, 241–250 (2018).

10. Magistretti, P. J. & Allaman, I. A cellular perspective on brain energy metabolism and functional imaging. Neuron 86, 883–901 (2015).

11. Hirata, K. et al. Altered brain energy metabolism related to astrocytes in Alzheimer’s disease. Ann. Neurol. 95, 104–115 (2024).

12. Mathys, H. et al. Single-cell multiregion dissection of Alzheimer’s disease. Nature 632, 858–868 (2024).

13. Gabitto, M. I. et al. Integrated multimodal cell atlas of Alzheimer’s disease. Nat. Neurosci. 27, 2366–2383 (2024).

14. Johnson, E. C. B. et al. Large-scale deep multi-layer analysis of Alzheimer’s disease brain reveals strong proteomic disease-related changes not observed at the RNA level. Nat. Neurosci. 25, 213–225 (2022).

15. Brindle, K. M. Imaging metabolism with hyperpolarized (13)C-labeled cell substrates. J. Am. Chem. Soc. 137, 6418–6427 (2015).

16. Sara, S. J. The locus coeruleus and noradrenergic modulation of cognition. Nat. Rev. Neurosci. 10, 211–223 (2009).

17. Ding, F. et al. α1-Adrenergic receptors mediate coordinated Ca2+ signaling of cortical astrocytes in awake, behaving mice. Cell Calcium 54, 387–394 (2013).

18. Dienel, G. A. & Cruz, N. F. Aerobic glycolysis during brain activation: adrenergic regulation and influence of norepinephrine on astrocytic metabolism. J. Neurochem. 138, 14–52 (2016).

19. Ludwig, C. et al. Data-independent acquisition-based SWATH-MS for quantitative proteomics: a tutorial. Mol. Syst. Biol. 14, e8126 (2018).

20. Lee, A. J. et al. Multi-region brain transcriptomes uncover two subtypes of aging individuals with differences in Alzheimer risk and the impact of APOEε4. iScience 29, 117038 (2026).

21. Vasilkovska, T. et al. Hyperpolarized ^13^C MRS can detect alterations in the astrocyte-neuron lactate shuttle (ANLS) in the healthy mouse brain. Proc. Intl. Soc. Mag. Reson. Med. 33, 3645 (2025).

22. Ono, M. et al. Altered astrocyte-neuron crosstalk in progressive supranuclear palsy: integrated evidence from proteomics and magnetic resonance spectroscopy. Acta Neuropathol. 151, 45 (2026).

23. Valla, J., Berndt, J. D. & Gonzalez-Lima, F. Energy hypometabolism in posterior cingulate cortex of Alzheimer’s patients: superficial laminar cytochrome oxidase associated with disease duration. J. Neurosci. 21, 4923–4930 (2001).

24. Liang, W. S. et al. Alzheimer’s disease is associated with reduced expression of energy metabolism genes in posterior cingulate neurons. Proc. Natl. Acad. Sci. U. S. A. 105, 4441–4446 (2008).

25. Sekar, S. et al. Alzheimer’s disease is associated with altered expression of genes involved in immune response and mitochondrial processes in astrocytes. Neurobiol. Aging 36, 583– 591 (2015).

26. Ono, M. et al. Deuteration enhances UV-induced hyperpolarization of [1-13C]pyruvate to trityl-level performance in vitro and in vivo. J. Cereb. Blood Flow Metab. 271678X261482014 (2026).

27. Vasilkovska, T. et al. Hyperpolarized 13C MRI uncovers early and progressive metabolic dysfunction in the hAPP-J20 mouse model of Alzheimer’s disease. Res. Sq. (2026) doi:10.21203/rs.3.rs-10437814/v1.

28. Romanov, N. et al. Disentangling genetic and environmental effects on the proteotypes of individuals. Cell 177, 1308–1318.e10 (2019).

29. Lebon, V. et al. Astroglial contribution to brain energy metabolism in humans revealed by 13C nuclear magnetic resonance spectroscopy: elucidation of the dominant pathway for neurotransmitter glutamate repletion and measurement of astrocytic oxidative metabolism. J. Neurosci. 22, 1523–1531 (2002).

30. Pellerin, L. & Magistretti, P. J. Glutamate uptake into astrocytes stimulates aerobic glycolysis: a mechanism coupling neuronal activity to glucose utilization. Proc. Natl. Acad. Sci. U. S. A. 91, 10625–10629 (1994).

31. Theofilas, P. et al. Locus coeruleus volume and cell population changes during Alzheimer’s disease progression: A stereological study in human postmortem brains with potential implication for early-stage biomarker discovery. Alzheimers. Dement. 13, 236– 246 (2017).

32. Zuend, M. et al. Arousal-induced cortical activity triggers lactate release from astrocytes. Nat. Metab. 2, 179–191 (2020).

33. Heneka, M. T. et al. Locus ceruleus controls Alzheimer’s disease pathology by modulating microglial functions through norepinephrine. Proc. Natl. Acad. Sci. U. S. A. 107, 6058–6063 (2010).

34. Prokopiou, P. C. et al. Astrocyte reactivity modifies the effects of locus coeruleus norepinephrine-related dysfunction on tau and its impact on cognition. Alzheimers. Dement. 22, e71703 (2026).

35. Braak, H. & Braak, E. Neuropathological stageing of Alzheimer-related changes. Acta Neuropathol. 82, 239–259 (1991).

36. Braak, H., Alafuzoff, I., Arzberger, T., Kretzschmar, H. & Del Tredici, K. Staging of Alzheimer disease-associated neurofibrillary pathology using paraffin sections and immunocytochemistry. Acta Neuropathol. 112, 389–404 (2006).

37. Montine, T. J. et al. National Institute on Aging-Alzheimer’s Association guidelines for the neuropathologic assessment of Alzheimer’s disease: a practical approach. Acta Neuropathol. 123, 1–11 (2012).

38. Nelson, P. T. et al. Limbic-predominant age-related TDP-43 encephalopathy (LATE): consensus working group report. Brain 142, 1503–1527 (2019).

39. Bennett, D. A. et al. Neuropathology of older persons without cognitive impairment from two community-based studies. Neurology 66, 1837–1844 (2006).

40. McKhann, G. et al. Clinical diagnosis of Alzheimer’s disease: report of the NINCDS-ADRDA Work Group under the auspices of Department of Health and Human Services Task Force on Alzheimer’s Disease. Neurology 34, 939–944 (1984).

41. Hyman, B. T. & Trojanowski, J. Q. Consensus recommendations for the postmortem diagnosis of Alzheimer disease from the National Institute on Aging and the Reagan Institute Working Group on diagnostic criteria for the neuropathological assessment of Alzheimer disease. J. Neuropathol. Exp. Neurol. 56, 1095–1097 (1997).

42. Percie du Sert, N., et al. The ARRIVE guidelines 2.0: Updated guidelines for reporting animal research. PLoS Biol. 18, e3000410 (2020).

43. Saito, T. et al. Single App knock-in mouse models of Alzheimer’s disease. Nat. Neurosci. 17, 661–663 (2014).

44. Santacruz, K. et al. Tau suppression in a neurodegenerative mouse model improves memory function. Science 309, 476–481 (2005).

45. Akai, M. et al. miR-PAIR: microRNA-protein analysis of integrative relationship for the identification of significantly working miRNAs. Biochim. Biophys. Acta Gen. Subj. 1869, 130746 (2025).

46. Tkác, I., Starcuk, Z., Choi, I. Y. & Gruetter, R. In vivo 1H NMR spectroscopy of rat brain at 1 ms echo time. Magn. Reson. Med. 41, 649–656 (1999).

47. Provencher, S. W. Estimation of metabolite concentrations from localized in vivo proton NMR spectra. Magn. Reson. Med. 30, 672–679 (1993).

48. Ono, M. et al. Awake brain MRSI reveals anesthetic sensitivity and regional aging effects on [13C]bicarbonate metabolism in mice. Front. Neuroimaging 4, 1506126 (2025).

49. Benjamini, Y. & Hochberg, Y. Controlling the false discovery rate: A practical and powerful approach to multiple testing. J. R. Stat. Soc. Series B Stat. Methodol. 57, 289– 300 (1995).

50. Love, M. I., Huber, W. & Anders, S. Moderated estimation of fold change and dispersion for RNA-seq data with DESeq2. Genome Biol. 15, 550 (2014).

51. Subramanian, A., et al. Gene set enrichment analysis: A knowledge-based approach for interpreting genome-wide expression profiles. Proc. Natl. Acad. Sci. U.S.A. 102, 15545–15550 (2005).

52. Wu, T. et al. clusterProfiler 4.0: A universal enrichment tool for interpreting omics data. Innovation (Camb.) 2, 100141 (2021).

53. Liberzon, A. et al. Molecular signatures database (MSigDB) 3.0. Bioinformatics 27, 1739–1740 (2011).

54. Hänzelmann, S., Castelo, R. & Guinney, J. GSVA: gene set variation analysis for microarray and RNA-seq data. BMC Bioinformatics 14, 7 (2013).

55. Ritchie, M. E. et al. limma powers differential expression analyses for RNA-sequencing and microarray studies. Nucleic Acids Res. 43, e47 (2015).

56. Russo, P. S. T. et al. CEMiTool: a Bioconductor package for performing comprehensive modular co-expression analyses. BMC Bioinformatics 19, 56 (2018).

57. Langfelder, P. & Horvath, S. WGCNA: an R package for weighted correlation network analysis. BMC Bioinformatics 9, 559 (2008).

58. Cahill, K. M., Huo, Z., Tseng, G. C., Logan, R. W. & Seney, M. L. Improved identification of concordant and discordant gene expression signatures using an updated rank-rank hypergeometric overlap approach. Sci. Rep. 8, 9588 (2018).

59. Hao, Y. et al. Dictionary learning for integrative, multimodal and scalable single-cell analysis. Nat. Biotechnol. 42, 293–304 (2024).

60. Hafemeister, C. & Satija, R. Normalization and variance stabilization of single-cell RNA-seq data using regularized negative binomial regression. Genome Biol. 20, 296 (2019).

61. Keren-Shaul, H. et al. A unique microglia type associated with restricting development of Alzheimer’s disease. Cell 169, 1276–1290.e17 (2017).

62. Habib, N. et al. Disease-associated astrocytes in Alzheimer’s disease and aging. Nat. Neurosci. 23, 701–706 (2020).

63. Choi, H. et al. Spatiotemporal characterization of glial cell activation in an Alzheimer’s disease model by spatially resolved transcriptomics. Exp. Mol. Med. 55, 2564–2575 (2023).

64. Korsunsky, I. et al. Fast, sensitive and accurate integration of single-cell data with Harmony. Nat. Methods 16, 1289–1296 (2019).

65. Okuda, S. et al. jPOST environment accelerates the reuse and reanalysis of public proteome mass spectrometry data. Nucleic Acids Res. 53, D462–D467 (2025).

