## Supplementary Figures 1-7 for "Glial involvement in mitochondrial metabolic alterations in Alzheimer’s disease"

[1] Institute for Quantum Life Science, National Institutes for Quantum Science and Technology, Chiba, Japan
[2] Department of Neurology and Neurological Science, Institute of Science Tokyo, Tokyo, Japan
[3] Department of Genetics and Genomic Sciences, Icahn School of Medicine at Mount Sinai, New York, NY, USA
[4] Nash Family Department of Neuroscience & Friedman Brain Institute, Icahn School of Medicine at Mount Sinai, New York, NY, USA
[5] Ronald M. Loeb Center for Alzheimer's Disease, Icahn School of Medicine at Mount Sinai, New York, NY, USA
[6] Icahn Institute for Data Science and Genomic Technology, Icahn School of Medicine at Mount Sinai, New York, NY, USA
[7] Estelle and Daniel Maggin Department of Neurology, Icahn School of Medicine at Mount Sinai, New York, NY, USA
[8] Graduate School of Biomedical and Health Sciences, Hiroshima University, Hiroshima, Japan
[9] Graduate School of Pharmaceutical Sciences, Tohoku University, Sendai, Japan
[10] Core for Spin Life Sciences, National Institute for Physiological Sciences, Okazaki, Japan
[11] Section of Brain Function Information, National Institute for Physiological Sciences, Okazaki, Japan
[12] Laboratory for Glia-Neuron Circuit Dynamics, RIKEN Center for Brain Science, Wako, Saitama, Japan
[13] Department of Life Science and Medical Bioscience, Graduate School of Advanced Science and Engineering, TWIns, Waseda University, Shinjuku-ku, Tokyo, Japan
[14] Department of Pathology, Brain Research Institute, Niigata University, Niigata, Japan
[15] Center for Human Brain Resource Initiative (ChBRI), Niigata University, Niigata, Japan
*These authors contributed equally: Kosei Hirata and Maiko Ono.

### Supplementary Information

Supplementary Fig. 1. Lower-magnification GFAP and NDUFA6 immunofluorescence in human PCC.

Supplementary Fig. 2. Co-expression network analysis of human PCC bulk RNA-seq identifies modules associated with cognition, neuropathology and glial signatures.

Supplementary Fig. 3. MRS-based assessment of additional hippocampal metabolites in APP-KI and wild-type mice.

Supplementary Fig. 4. MRS-based metabolite assessment in the cortex of APP-KI and wild-type mice.

Supplementary Fig. 5. MRS-based metabolite assessment in the hippocampus of rTg4510 and non-Tg mice.

Supplementary Fig. 6. Regional distributions of Visium v2 module scores in WT and APP-KI mice.

Supplementary Fig. 7. Slab placement for MRSI.

Supplementary Tables 1–13 are provided as a separate Excel workbook.

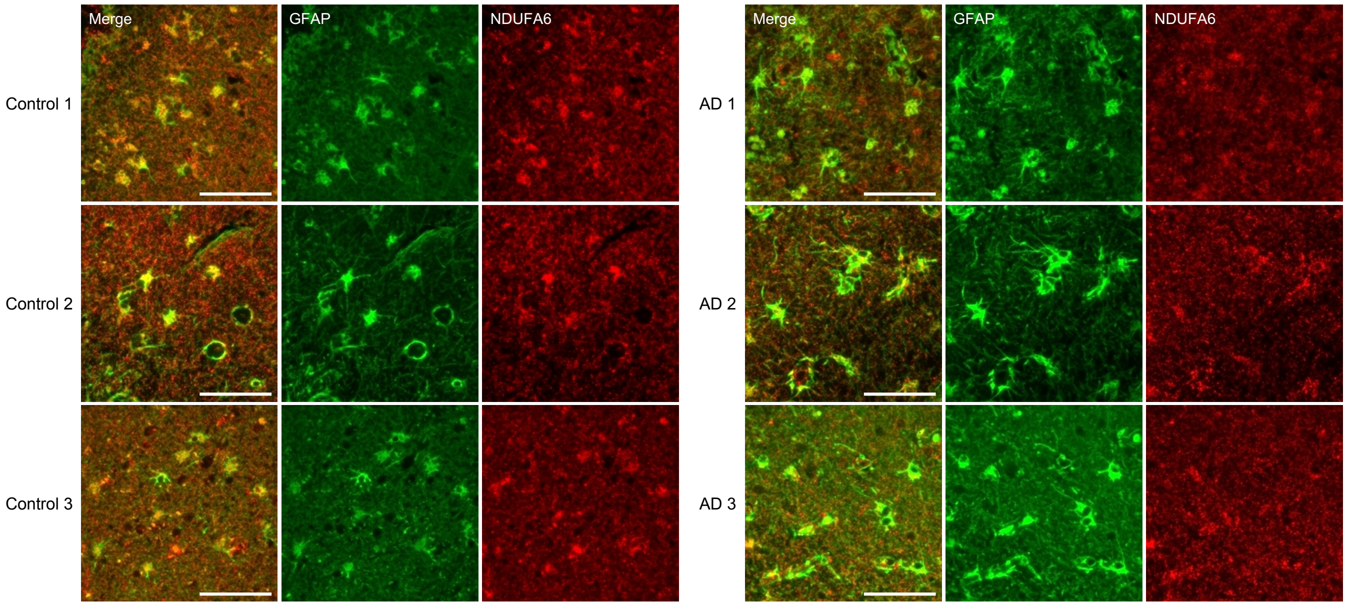


**Supplementary Fig. 1. Lower-magnification GFAP and NDUFA6 immunofluorescence in human PCC.**

Lower-magnification images of the merged channels, GFAP (green) and NDUFA6 (red), from left to right within each group, from the same three control and three AD cases shown in Fig. 1h. Rows show individual cases within each diagnostic group; equal row numbers do not indicate matched pairs. The images provide tissue context for the qualitative observations in Fig. 1h and were not used for quantitative comparison between diagnostic groups. Scale bars, 50 μm.

Abbreviations: GFAP, glial fibrillary acidic protein; PCC, posterior cingulate cortex; AD, Alzheimer's disease.

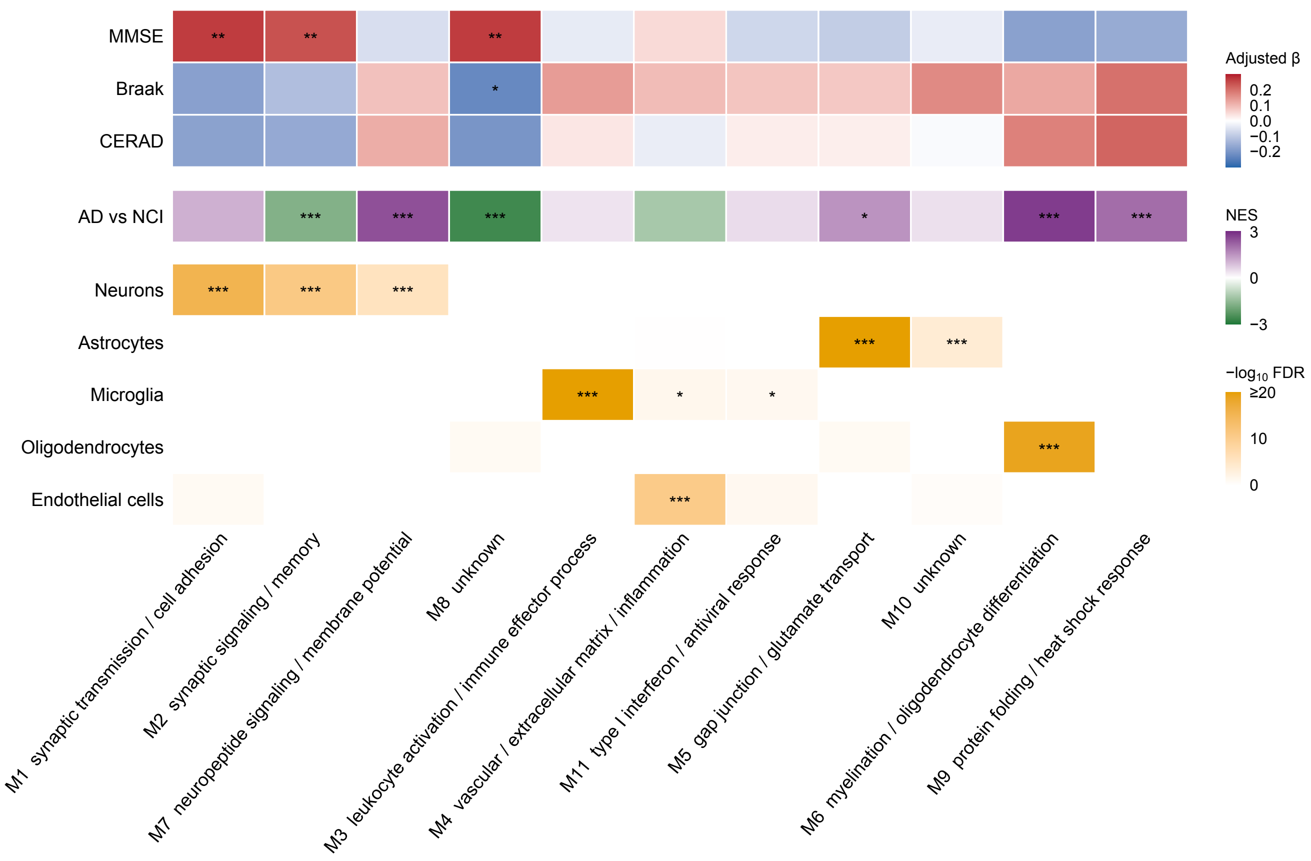


**Supplementary Fig. 2. Co-expression network analysis of human PCC bulk RNA-seq identifies modules associated with cognition, neuropathology and glial signatures.**

Summary of 11 co-expression modules identified by CEMiTool in AD ($n$ = 117) and NCI ($n$ = 69) PCC samples. Columns represent modules, labelled with their functional annotations. Top, covariate-adjusted standardized regression coefficients (β) for associations between module eigengenes and MMSE score, Braak stage or CERAD score. Red and blue indicate positive and negative associations, respectively. Middle, normalized enrichment scores (NES) for module activity in AD relative to NCI; purple indicates higher activity in AD and green indicates lower activity. Bottom, enrichment of cell-type marker genes within each module, assessed using one-sided Fisher's exact tests with Benjamini–Hochberg correction within each cell type. Colour intensity indicates $-\log_{10}$ FDR, capped at 20. Across all sections, asterisks indicate *FDR < 0.05, **FDR < 0.01 and ***FDR < 0.001.

Abbreviations: AD, Alzheimer's disease; PCC, posterior cingulate cortex; NCI, no cognitive impairment; MMSE, Mini-Mental State Examination; CERAD, Consortium to Establish a Registry for Alzheimer's Disease; FDR, false discovery rate.

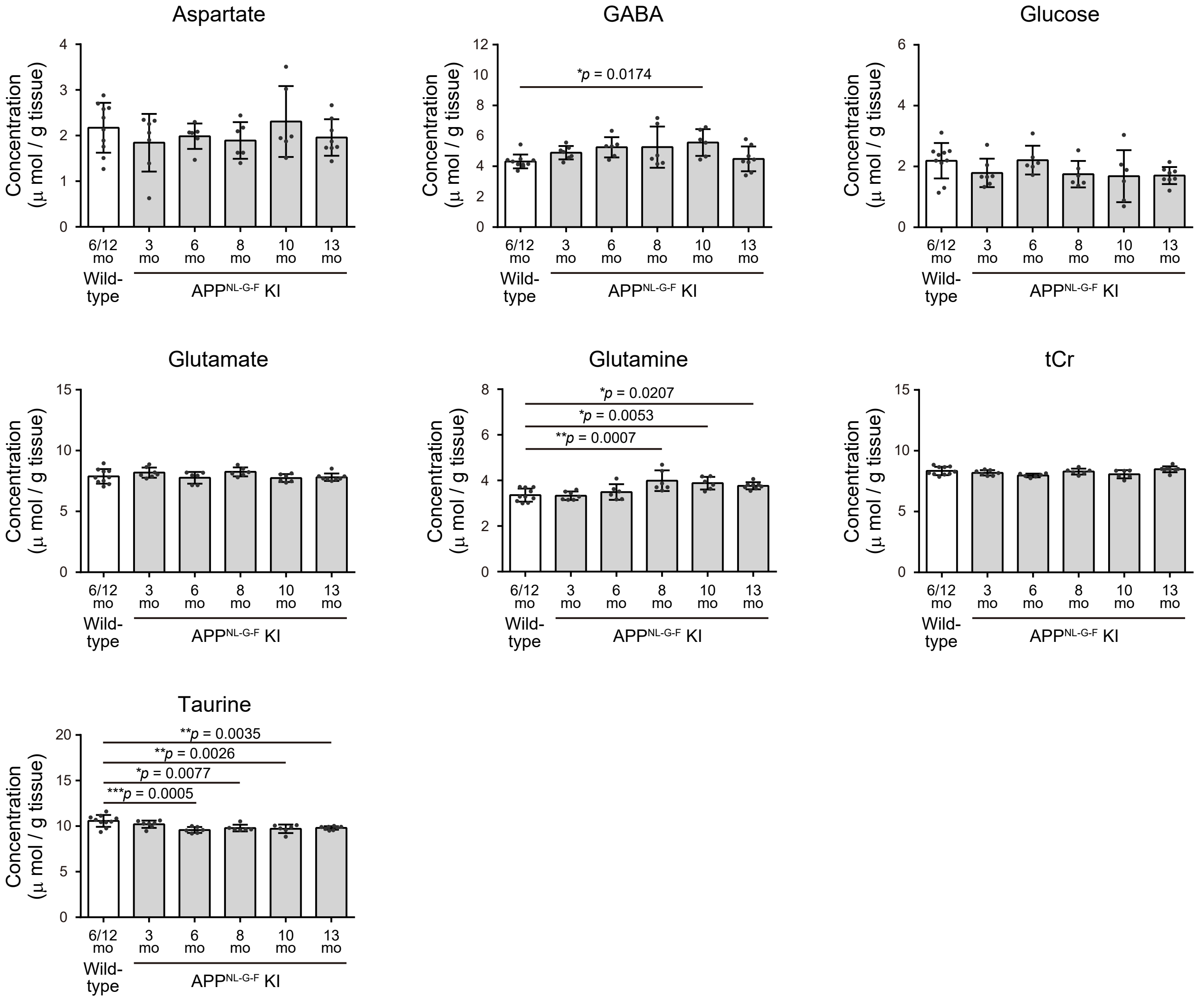


**Supplementary Fig. 3. MRS-based assessment of additional hippocampal metabolites in APP-KI and wild-type mice.**

Hippocampal concentrations of aspartate, GABA, glucose, glutamate, glutamine, tCr, and taurine in wild-type mice (6 and 12 months) and APP-KI mice (3, 6, 8, 10, and 13 months). Data are shown as individual values with mean ± standard deviation. **P* < 0.05, ***P* < 0.005, ****P* < 0.0005 by one-way ANOVA followed by Dunnett's multiple comparisons test.

Abbreviations: APP-KI, *App*$^{NL\text{-}G\text{-}F}$ knock-in; GABA, γ-aminobutyric acid; tCr, total creatine (creatine + phosphocreatine); ANOVA, analysis of variance; MRS, magnetic resonance spectroscopy.

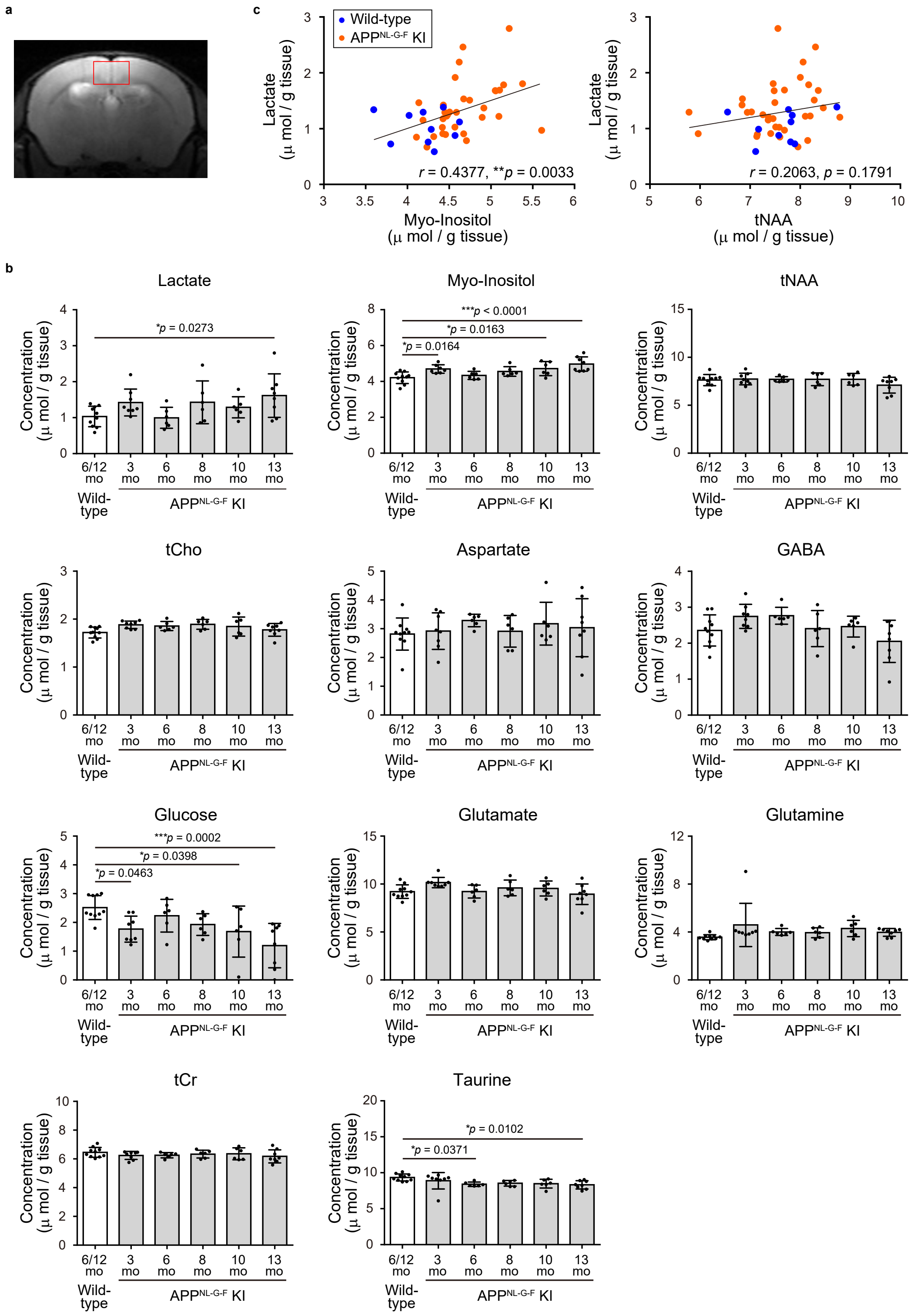


**Supplementary Fig. 4. MRS-based metabolite assessment in the cortex of APP-KI and wild-type mice.**

(**a**) Representative single-voxel placement (red squares) in the cortex (1.2 × 2.5 × 3.0 $mm^3$) overlaid on a coronal $T_2$-weighted MR image. (**b**) Cortical concentrations of lactate, *myo*-inositol, tNAA, tCho, aspartate, GABA, glucose, glutamate, glutamine, tCr, and taurine in wild-type mice (6 and 12 months) and APP-KI mice (3, 6, 8, 10, and 13 months). Data are shown as individual values with mean ± standard deviation. $^*P < 0.05$, $^{***}P < 0.0005$ by one-way ANOVA followed by Dunnett's multiple comparisons test. (**c**) Correlations between cortical metabolite concentrations, specifically lactate and *myo*-inositol (left), and lactate and tNAA (right), across all animals. $^{**}P < 0.005$ using Pearson's correlation analysis.

Abbreviations: APP-KI, $App^{NL\text{-}G\text{-}F}$ knock-in; MRS, magnetic resonance spectroscopy; MR, magnetic resonance; tNAA, total N-acetylaspartate (NAA + NAAG); NAA, N-acetylaspartate; NAAG, N-acetylaspartylglutamate; tCho, total choline (glycerophosphocholine + phosphocholine); GABA, γ-aminobutyric acid; tCr, total creatine (creatine + phosphocreatine); ANOVA, analysis of variance.

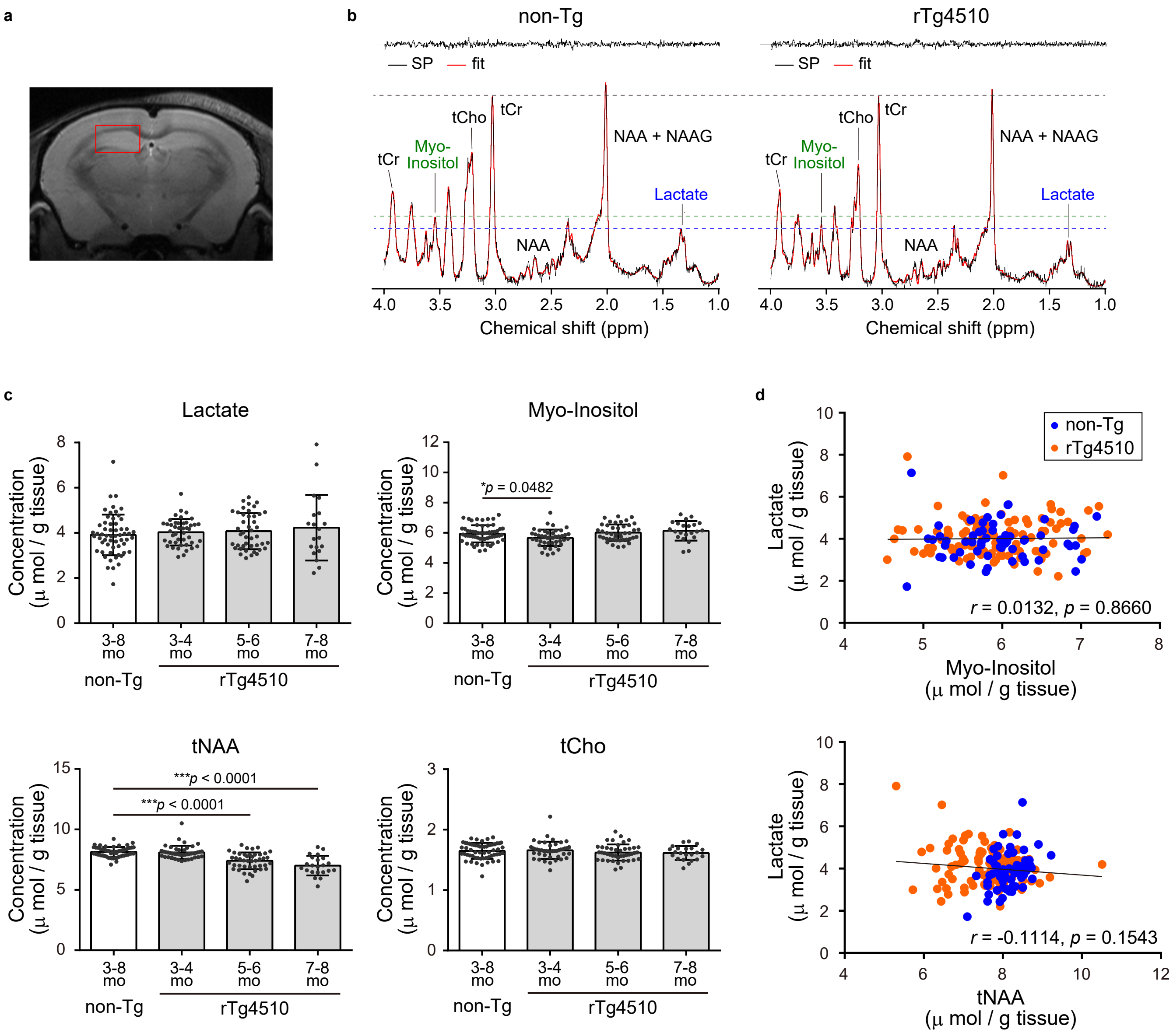


**Supplementary Fig. 5. MRS-based metabolite assessment in the hippocampus of rTg4510 and non-Tg mice.**

(**a**) Representative single-voxel placement (red squares) in the hippocampus (1.2 × 2.0 × 2.2 $mm^3$) overlaid on a coronal $T_2$-weighted MR image. (**b**) Representative single-voxel $^1$H-MRS (PRESS) spectra and corresponding fits from 7–8-month-old non-Tg and rTg4510 mice. tCr = creatine + phosphocreatine, tCho = glycerophosphocholine + phosphocholine, NAA = N-acetylaspartate, NAAG = N-acetylaspartylglutamate. (**c**) Hippocampal concentrations of lactate, myo-inositol, tNAA, and tCho in non-Tg mice (3–8 months) and rTg4510 mice (3–4, 5–6, and 7–8 months). Data are shown as individual values with mean ± standard deviation. $*P < 0.05$, $***P < 0.0005$ by one-way ANOVA followed by Dunnett's multiple comparisons test. (**d**) Correlations between hippocampal metabolite concentrations, specifically lactate and myo-inositol (upper), and lactate and tNAA (lower), across all animals, assessed using the Pearson correlation coefficient.

Abbreviations: non-Tg, non-transgenic; MRS, magnetic resonance spectroscopy; MR, magnetic resonance; $^1$H-MRS, proton magnetic resonance spectroscopy; PRESS, point-resolved spectroscopy; tCr, total creatine (creatine + phosphocreatine); tCho, total choline (glycerophosphocholine + phosphocholine); tNAA, total N-acetylaspartate (NAA + NAAG); NAA, N-acetylaspartate; NAAG, N-acetylaspartylglutamate; ANOVA, analysis of variance.

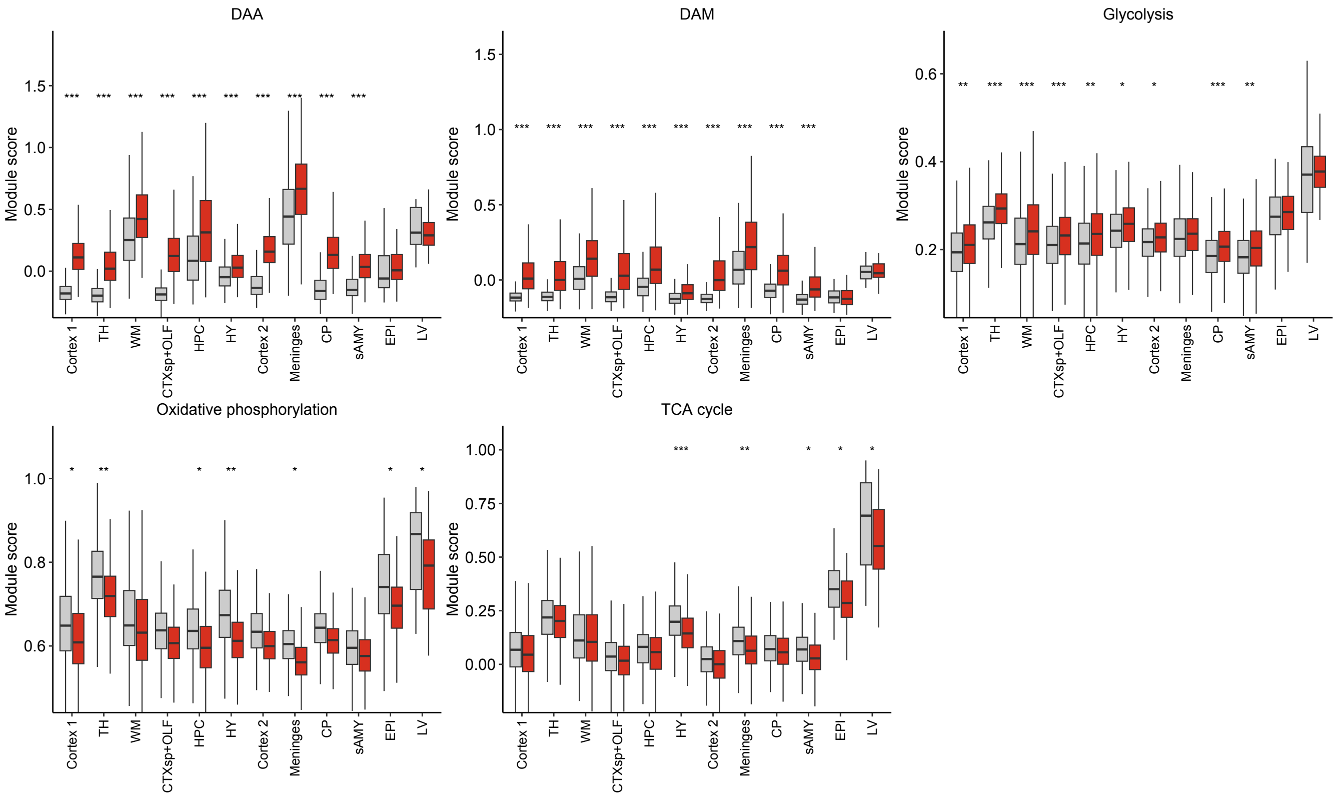


**Supplementary Fig. 6. Regional distributions of Visium v2 module scores in WT and APP-KI mice.**

Box plots of spot-level DAA, DAM, glycolysis, oxidative phosphorylation and TCA-cycle module scores in each of the 12 anatomical regions (n = 4 mice per group; grey, WT; red, APP-KI). Boxes show the median and interquartile range, and whiskers extend to 1.5 times the interquartile range; outlying spots are not shown. Asterisks indicate the FDR of the linear mixed models summarized in Fig. 5c (*FDR < 0.05, **FDR < 0.01 and ***FDR < 0.001).

Abbreviations: DAA, disease-associated astrocyte; DAM, disease-associated microglia; TCA, tricarboxylic acid; TH, thalamus; WM, white matter; CTXsp+OLF, cortical subplate and olfactory areas; HPC, hippocampus; HY, hypothalamus; CP, caudoputamen; sAMY, striatum-like amygdalar nuclei; EPI, epithalamus; LV, lateral ventricle; WT, wild type; APP-KI, AppNL-G-F knock-in; FDR, false discovery rate.

a

b

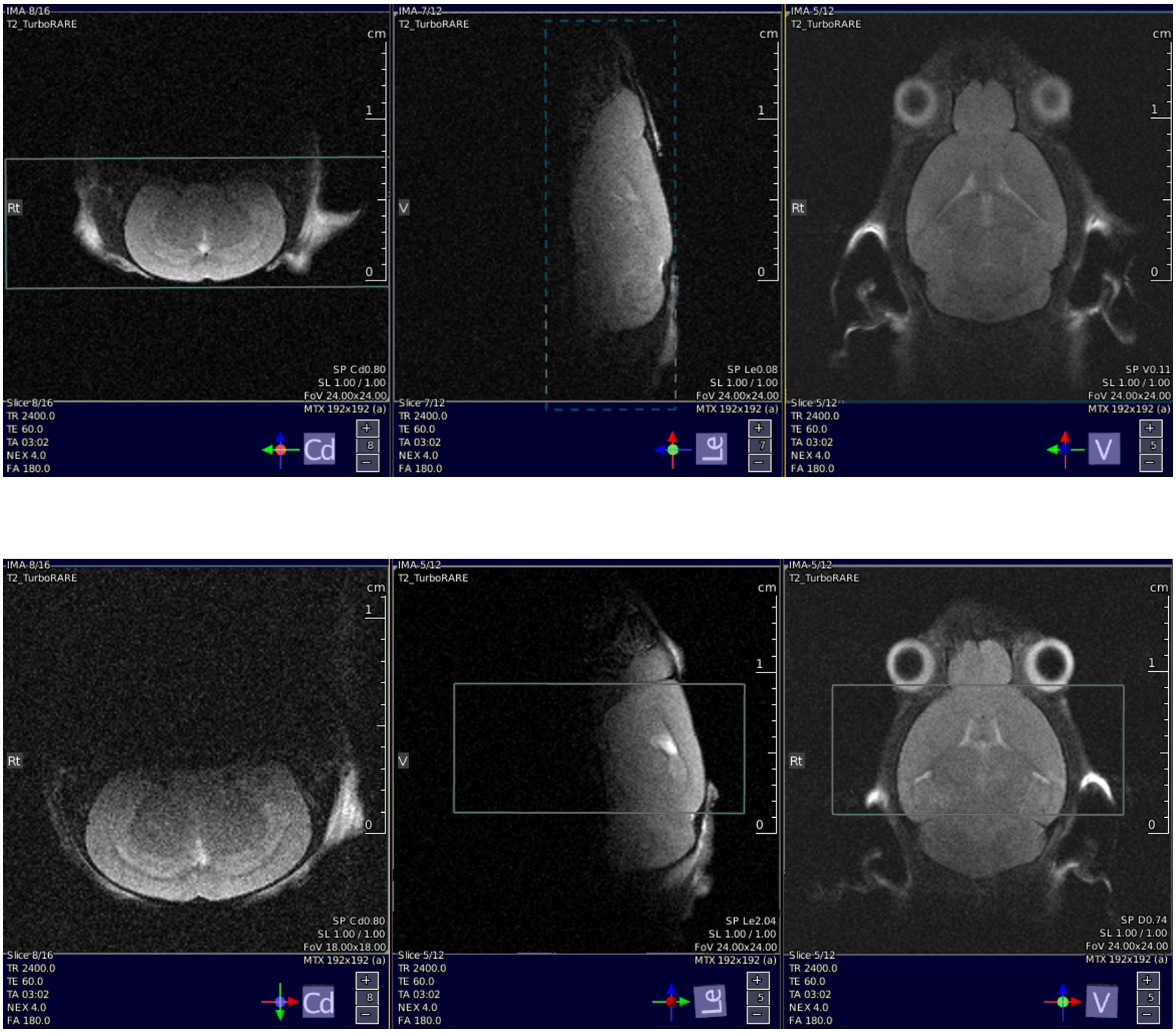


**Supplementary Fig. 7. Slab placement for MRSI.**

(a, b) Representative MRSI slabs positioned in the coronal plane to cover the whole brain (**a**) or in the axial plane to primarily cover the hippocampus and cortex (**b**).

Abbreviation: MRSI, magnetic resonance spectroscopic imaging.
